# The biological functions of Naa10 – from amino-terminal acetylation to human disease

**DOI:** 10.1101/014324

**Authors:** Max Doerfel, Gholson J. Lyon

## Abstract

N-terminal acetylation (NTA) is one of the most abundant protein modifications known, and the N-terminal acetyltransferase (NAT) machinery is conserved throughout all Eukarya. Over the past 50 years, the function of NTA has begun to be slowly elucidated, and this includes the modulation of protein-protein interaction, protein-stability, protein function, and protein targeting to specific cellular compartments. Many of these functions have been studied in the context of Naa10/NatA; however, we are only starting to really understand the full complexity of this picture. Roughly, about 40 % of all human proteins are substrates of Naa10 and the impact of this modification has only been studied for a few of them. Besides acting as a NAT in the NatA complex, recently other functions have been linked to Naa10, including post-translational NTA, lysine acetylation, and NAT/KAT-independent functions. Also, recent publications have linked mutations in Naa10 to various diseases, emphasizing the importance of Naa10 research in humans. The recent design and synthesis of the first bisubstrate inhibitors that potently and selectively inhibit the NatA/Naa10 complex, monomeric Naa10, and hNaa50 further increases the toolset to analyze Naa10 function.

## Introduction

N^α^-terminal acetylation (NTA) is one of the most abundant modifications of eukaryotic proteins. Today it is believed that the majority of the proteome of higher organisms is fully or partially acetylated. In fact, recent large-scale proteomics analyses have identified peptides that were fully or partially acetylated at their designated N-terminus in the following percentages: 13-19% in *Halobacterium salinarum* and *Natronomonas pharaonis* (Falb et al., 2006; Aivaliotis et al., 2007), 29% in *Haloferax volcanii* (Kirkland et al., 2008), about 16% in 45 tested bacteria ((Bonissone et al., 2013), 60-70% in *S. cerevisiae* (Arnesen et al., 2009b; Van Damme et al., 2011c; Bonissone et al., 2013; Van Damme et al., 2014), 75% in *Drosophila melanogaster* (Goetze et al., 2009), 90% in *Arabidopsis thaliana* (Bienvenut et al., 2012), at least 4% in *C. elegans* (Mawuenyega et al., 2003), 83% in mouse (Lange and Overall, 2011), 90% in human erythrocytes (Lange et al., 2014) and 85% in HeLa cells (Arnesen et al., 2009b; Van Damme et al., 2011c). However, these values do not necessarily reflect the whole proteomes. A recent computational analysis of large-scale proteome analyses was used to develop a prediction software for NTA in archae (*P. furiosus*, *T. acidophilum*, *H. salinarum* and *N. pharaonis*), animals (*Homo sapiens*, *Caenorhadbitis elegans*, and *Drosophila melanogaster)*, plants (*A. thaliana* and *Oryza sativa*) and fungi (*S. cerevisiae* and *N. crassa*). The analysis revealed a bias for N-terminal acetylated proteins in highly abundant cytosolic proteins (Martinez et al., 2008). This bias could indicate that the reported percentage of acetylation is higher than the percentage in the actual proteomes: archeae 1-6.5%, animal 58 %, fungi and plants 60% (Martinez et al., 2008). Furthermore, in some studies only annotated N-termini were analyzed, others included N-termini derived from alternative translation initiation sites.

*In vitro* data suggests that NTA occurs mainly co-translationally on the emerging polypeptide chain at a length of approximately 25-80 residues (Strous et al., 1973; Filner and Marcus, 1974; Strous et al., 1974; Driessen et al., 1985; Gautschi et al., 2003), either on the initiating methionine (iMet) or on the second amino acid after methionine cleavage, also known as N-terminal methionine excision (NME) (Kendall and Bradshaw, 1992; Xiao et al., 2010; Bonissone et al., 2013). The removal of the iMet is the first occurring widespread protein modification and involves peptide deformylases and methionine aminopeptidases (MetAPs) (Giglione et al., 2014). In addition to co-translational acetylation, accumulating evidence also supports the occurrence of post-translational N^α^-acetylation. For example, the ribosomal protein L7/L12 in *E. coli* becomes acetylated post-translationally depending on the availability of nutrients (Gordiyenko et al., 2008). Furthermore, proteomic analyses identified NTA of internal peptides, further supporting the idea of post-translational acetylation (Helbig et al., 2010; Helsens et al., 2011). This is especially interesting for many proteins that are imported into organelles, after which the cleaved mature N-terminus of the protein (now missing its target/transit peptide) is acetylated by dedicated NATs that reside in the respective target organelle as shown for yeast mitochondrial localized proteins (Van Damme et al., 2014) or chloroplast proteins in *Chlamydomonas reinhardtii* and *Arabidopsis thaliana* (Zybailov et al., 2008; Bienvenut et al., 2011; Bienvenut et al., 2012).

NTA is catalyzed by distinct N^α^-acetyltransferases (NATs) that belong to the GCN5-related N-acetyltransferase (GNAT) family, a diverse family that catalyze the transfer of an acetyl group from acetyl-CoA to the primary amine of a wide variety of substrates from small molecules to large proteins (Vetting et al., 2005). Besides the NATs, this protein family also includes/contains lysine acetyltransferases (KATs) and histone acetyltransferases (HATs) (Marmorstein and Zhou, 2014).

In 2009, a new nomenclature for the N^α^-acetyltransferases was introduced (Polevoda et al., 2009), in which the concept of multi-protein complexes for NATs was formalized. In humans, six NATs, NatA-F, were defined that specifically co-translationally catalyze the acetylation of the N^α^-terminal amino group of a well-defined subset of proteins, although N^ε^-acetylation of internal lysines has also been reported (Kalvik and Arnesen, 2013). NatA consists of the catalytic subunit Naa10 and the auxiliary subunit Naa15 and acetylates small side chains such as Ser, Ala, Thr, Gly, Val and Cys after the initiator methionine has been cleaved by methionine aminopeptidases (via NME) (see Figure 1). NatB and NatC are defined as multimeric complexes containing the catalytic subunits Naa20 and Naa30 and the auxiliary subunits Naa25 and Naa35/Naa38, respectively. They acetylate proteins with their methionine retained. The only known substrates for NatD (Naa40) are histone H2A and H4. Naa50 is the catalytic subunit of NatE, with a substrate specificity for N-termini starting with methionine followed by Leu, Lys, Ala and Met (Van Damme et al., 2011b). NatF is composed of Naa60 and has a substrate specificity that partially overlaps with NatC and NatE. It is important to note that this might not be the complete picture, as there are possibly other proteins binding and interacting with proteins in these NATs as currently defined [for reviews see (Arnesen, 2011; Van Damme et al., 2011a; Starheim et al., 2012; Aksnes et al., 2015a)]

**Figure 1:**
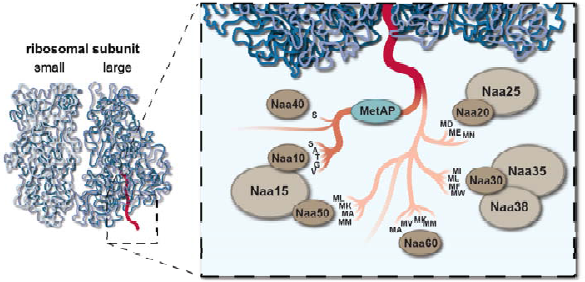
The co-translational N-terminal protein modification process. As soon as the nascent polypeptide chain emerges from the ribosome exit tunnel, the initiator methionine is cleaved by methionine aminopeptideases (MetAPs) if the following amino acid is small and uncharged. For the sake of simplicity, this process is illustrated by one enzyme despite the fact that other enzymes including peptide deformylases are involved, depending on organism and cellular cormpartment. Subsequently, the new N-termini can get acetylated by NatA, composed of the catalytic Naa10 and the auxiliary subunit Naa15. The majority of cytosolic proteins fall into this category. If the iMet is not processed, NTA can be accomplished by NatB (composed of Naa20 and Naa25), NatC (Naa30, Naa35, Naa38), NatD (Naa40) NatE (Naa50 and possibly Naa15) and NatF (Naa60). Figure modified from (Kalvik and Arnesen, 2013).

### 1.1 General Functions of Amino-terminal acetylation

Despite its discovery more than 50 years ago, very little is known about the biological function of NTA. For many years, it has been generalized from a few examples that NTA broadly protects many proteins from degradation. This was supported by the fact that acetylation of globin and lysozyme prevents their degradation by the ubiquitin proteolytic system from reticulocytes (Hershko et al., 1984). In line with this, the NTA of enkephalins diminishes their proteolytic cleavage by aminopeptidase M (Jayawardene and Dass, 1999), improves the chemical stability of guinea pig myelin basic protein (de Haan et al., 2004) and protects glucagon-like peptide (GLP-1) from dipeptidyl peptidase IV (DPP-IV)-mediated degradation (John et al., 2008). Analysis of the half-life of β-galactosidase in a split-ubiquitin system showed that proteins having N-terminal amino acids that are prone to acetylation (Met, Ser, Ala, Thr, Val, or Gly) have a relatively long half-life, whereas Arg, Lys, Phe, Leu, or Asp at the amino-terminus have very short half-lives (Bachmair et al., 1986). One way that NTA could contribute to protein stability is by blocking the access for N-terminal ubiquitination as shown for p16 and p14/p19^ARF^ (Ben-Saadon et al., 2004; Ciechanover and Ben-Saadon, 2004; Kuo et al., 2004). Also, p21^Cip1^ is acetylated (most likely by NatA) at its N-terminus, whereas an N-terminally tagged variant that abolishes N^α^-acetylation becomes ubiquitylated (Chen et al., 2004). However, other studies suggest that NTA has no effects on protein stability (Greenfield et al., 1994; Yi et al., 2011) and recent studies even showed that NTA might rather promote protein ubiquitination and degradation, depending on the cellular availability of interaction partners. The ubiquitin ligases Doa10 and Not4 recognize Nt-acetylated Ala, Val, Ser, Thr, and Cys and earmark acetylated substrates for degradation (Hwang et al., 2010; Shemorry et al., 2013). NTA therefore creates protein degradation signals (^Ac^N-degrons) that are targeted by the Ac/N-end rule pathway resulting in ubiquitylation and proteasome-mediated degradation by the Doa10 E3 N-recognin, in conjunction with the Ubc6 and Ubc7 E2 enzymes (Varshavsky, 2011). Conversely, multiple Doa10 substrates do not require N^α^-acetylation for their degradation, and acetylation has only mild effects on the stability of the tested substrates (Zattas et al., 2013). These discrepancies can be explained with the findings that N^α^-acetylation can have stabilizing effects when an interaction partner is involved. In this case, the acetylated N-termini recruits the interaction partners that then shield the ^Ac^N-degron, preventing ubiquitinylation and degradation to regulate subunit stoichiometries (Shemorry et al., 2013; Park et al., 2015). In agreement with this, it is widely accepted that N^α^-terminal acetylation can act as an avidity enhancer within protein complexes (Deakin et al., 1980; Scott et al., 2011; Nazmi et al., 2012). In this context, since the recent identification of an E2 ubiquitin ligase Ube2w that targets the α-amino group of proteins, it was speculated that – in this case – NTA could protect proteins from being proteasomally degraded by blocking of the target site (Aksnes et al., 2015a). NTA may also regulate NEDDylation, since N^α^-terminal acetylation of the E2 enzymes UBC12 and UBE2F has been found to be required for recognition by DCN-like co-E3s which promote ligation of NEDD8 to cullin targets (Scott et al., 2011; Monda et al., 2013; Scott et al., 2014). This is also another good example where NTA acts as an avidity enhancer.

Other studies suggest that NTA may play a role in the structural stabilization of N-terminally flexible proteins, as bioinformatics analysis of the yeast proteome showed that proteins with N-terminal-disordered regions are more likely to be acetylated (Holmes et al., 2014). In line with that, many studies have shown that NTA can stabilize an N-terminal α-helix (Shoemaker et al., 1987; Fairman et al., 1989; Chakrabartty et al., 1993; Doig et al., 1994; Greenfield et al., 1994; Jarvis et al., 1995; Fauvet et al., 2012; Kang et al., 2012; Kang et al., 2013).

Another function of NTA has been implicated in protein sorting and secretory processes. In yeast, the ARF-like GTPase Arl3p/ARP is acetylated by NatC and this modification is required for its targeting to the Golgi apparatus, possibly through the acetylation-dependent interaction with the integral membrane protein Sys1p (Setty et al., 2004). Simultaneously, a different group confirmed that Sys1p is the receptor for Arl3p and knockout of NatC or mutation of the NatC complex to abrogate its acetyltransferase activity resulted in failure to target Arl3p to the Golgi (Behnia et al., 2004). Furthermore, targeting of the human homologue of Arl3p, ARFRP1, is dependent on Sys1p and mutation of the N-terminus of ARFRP1, that would abrogate acetylation by NatC, induced its mis-localization in COS cells (Behnia et al., 2004). This and the fact that the N-terminus of Arl3p is a potential NatC substrate in *S. cerevisiae*, *D. melanogaster*, *C. elegans* and plants indicates that this system is well conserved. Two other human lysosomal Arf-like GTPases, Arl8a and Arl8b (also known as Arl10b/c and Gie1/2), and their single homologue in *Drosophila* are potential substrates of NatC, and mass spectrometric analyses confirmed that human Arl8b is N-terminally acetylated (Hofmann and Munro, 2006). Later *in vitro* acetylation assays showed that Arl8b is acetylated by NatC and knockdown of the catalytic subunit of NatC (Starheim et al., 2009a) or replacement of the leucine in position 2 with alanine (Hofmann and Munro, 2006) resulted in a loss of its lysosomal localization. It should be mentioned that the protein was still found to be acetylated, presumably by NatA following removal of the initiator methionine (Hofmann and Munro, 2006), indicating that specifically the acetylated methionine rather than acetylation itself is important for lysosomal targeting of Arl8b. Also, the inner nuclear membrane protein Trm1-II was found to be mislocalized to the nucleoplasm, when NatC was knocked out or when the penultimate amino acid was mutated to inhibit NatC-dependent NTA (Murthi and Hopper, 2005). On the other hand, systematic analysis of predicted N-terminal processing in yeast showed that cytoplasmic proteins are typically acetylated, whereas those destined for secretion via the ER are largely unmodified (Forte et al., 2011). Mutation of the N-terminal amino acid of the secretory protein carboxypeptidase Y, which allowed acetylation of this protein, inhibited targeting to the ER (Forte et al., 2011). However, fluorescence microscopy analysis in yeast indicated unaltered subcellular localization patterns for all 13 studied NatC substrates, after disruption of the NatC catalytic subunit (Aksnes et al., 2013). Furthermore, no disruption of the nuclear membrane, endoplasmic reticulum, Golgi apparatus, mitochondria, or bud neck was observed upon NatC deletion, suggesting the intactness of these organelles and subcellular structures as judged by the unchanged shape, number, size and distribution in the cell (Aksnes et al., 2013). A follow-up study showed that not NatC but rather NatF (Naa60) is associated with Golgi membranes and the ER, and disruption of this Naa60 induces Golgi fragmentation (Aksnes et al., 2015b). Taken together, this indicates that NatC is not – at least not in general – a determinant for substrate subcellular localization (Aksnes et al., 2013). Similarly, fluorescence microscopy analysis of 13 NatB substrates in wild type and *NAA20*Δ yeast cells revealed that acetylation by NatB is not a general signal for protein localization (Caesar et al., 2006).

Aside from the above, NTA has been shown to affect protein function and/or activity in a variety of cases, including hemoglobin isoforms (Scheepens et al., 1995; Ashiuchi et al., 2005), phospholamban (PLB) (Starling et al., 1996), N-TIMPs (N-terminal inhibitory domains of TIMPs /inhibitors of metalloproteinases (Van Doren et al., 2008), parvalbumin (Permyakov et al., 2012), melanocyte-stimulating hormone (MSH) in the barfin flounder (*Verasper moseri*) (Kobayashi et al., 2009), the contractile proteins actin and tropomyosin in fission and budding yeast (Polevoda et al., 2003; Singer and Shaw, 2003; Coulton et al., 2010) as well as the stress-induced carboxypeptidase Y inhibitor Tfs1p in yeast (Caesar and Blomberg, 2004).

In addition, NTA has been linked to various diseases, including apoptosis and cancer (Kalvik and Arnesen, 2013), host parasite interaction in malaria (Chang et al., 2008), and has been discussed to play a role in Parkinson’s disease (see below). As pointed out in earlier reviews: “Although…[NTA]…is essential for cell viability and survival, very little is known about the physiological reasons associated with this crucial role” (Giglione et al., 2014) and “there may be a variety of acetylation-dependent functions depending on the target protein, rather than one general function [and] there is even the possibility that this modification affects the function of only very few proteins” (Arnesen, 2011). The cellular phenotypes observed by disruption of the different NATs has been summarized in a very recent review (Aksnes et al., 2015a).

The best studied N^α^-acetyltransferase NatA consists of the catalytic subunit Naa10 and the auxiliary subunit Naa15. In this review we mainly concentrate on Naa10 structure and function and discuss recent developments in the field.

### 1.2 The NatA complex

As mentioned above, the NatA complex consists at least of the auxiliary and catalytic subunit, Naa15 and Naa10, respectively and is evolutionarily conserved from yeast to vertebrates (Mullen et al., 1989; Park and Szostak, 1992; Sugiura et al., 2003; Arnesen et al., 2005a). We adopt here the nomenclature of inserting letters to indicate the species about which we are discussing, so yNatA refers to NatA in yeast, where we are specifically referring to *S. cerevesiae*, hNatA refers to NatA in humans, and mNatA refers to NatA in mice. However, this nomenclature in 2009 did not address other species, and it might be worth updating the nomenclature at some future international meeting focused on the NATs.

In any case, there is good *in vitro* and *in vivo* evidence that yNatA acetylates the N-termini of small side chains like serine, alanine, glycine and threonine (Arnold et al., 1999; Polevoda et al., 1999) and NatA from humans has identical or nearly identical specificities, acetylating proteins starting with small side chains like serine, glycine, alanine, threonine and cysteine (Arnesen et al., 2009b; Van Damme et al., 2011b; Van Damme et al., 2011c), after the removal of the initiator methionine by methionine aminopeptidases. It was found that heterologous combinations of human and yeast Naa10p and Naa15p are not functional in yeast, suggesting significant structural subunit differences between the proteins from the different species (Arnesen et al., 2009b). (Met-)Ala-N-termini are more prevalent in the human proteome, whereas (Met)-Ser-N-termini are more abundant in the yeast proteome (Van Damme et al., 2011c). Accordingly, hNatA displays a preference towards these Ala-N-termini whereas yNatA seems to be the more efficient in acetylating Ser-starting N-termini, indicating that NatA substrate specificity/efficiency of Nt-acetylation has co-evolved with the repertoire of NatA type substrates expressed (Van Damme et al., 2014).

Size-exclusion chromatography and circular dichroism showed that purified human Naa10 consists of a compact globular region comprising two thirds of the protein and a flexible unstructured C-terminus (Sánchez-Puig and Fersht, 2006). The recent X-ray crystal structure of the 100 kD holo-NatA (Naa10/Naa15) complex from *S. pombe* revealed that the auxiliary subunit Naa15 is composed of 37 α-helices ranging from 8 to 32 residues in length, among which 13 conserved helical bundle tetratricopeptide repeat (TPR) motifs can be identified (Liszczak et al., 2013). These Naa15 helices form a ring-like structure that wraps completely around the Naa10 catalytic subunit (Liszczak et al., 2013). TPR motifs mediate protein-protein interactions, and it was speculated that TPR might be important for interaction with other NatA-binding partners such as the ribosome, Naa50/NatE and the HYPK chaperone (Liszczak et al., 2013), but this needs to be proven in future experiments. We discuss the possible interaction with NatE in more detail below. Naa10 adopts a typical GNAT fold containing a N-terminal α1–loop–α2 segment that features one large hydrophibic interface and exhibits the most intimate interactions with Naa15, a central acetyl CoA-binding region and C-terminal segments that are similar to the corresponding regions in Naa50 (Liszczak et al., 2013). The X-ray crystal structure of archaeal *T. volcanium* Naa10 has also been reported, revealing multiple distinct modes of acetyl-Co binding involving the loops between β4 and α3 including the P-loop (Ma et al., 2014). To our knowledge, there is not yet any published cryo-electron microscopy data regarding larger complexes between the ribosome, nascent polypeptide chain and any NATs.

Besides acting in a complex, it has been shown that a fraction of human Naa10 exists independent of Naa15 in the cytoplasm and is able to acetylate acidic side chains like aspartate and glutamate in γ- and β-actin (Van Damme et al., 2011b; Foyn et al., 2013a). These Type I actins are natural NatB substrates (iMet followed by an amino acid with an acidic side chain) and are therefore initially acetylated by NatB in yeast and humans (Van Damme et al., 2012). However, further processing/cleavage by an N^α^-acetylaminopeptidase (ANAP), which specifically removes the N-terminal Ac-Met or Ac-Cys from actin exposes the acidic N-terminal residue (Polevoda and Sherman, 2003b), which can be subsequently acetylated by Naa10. This substrate switching of Naa10 from small side chains towards acidic side chains could be explained by comparing the X-ray crystal structures of complexed (Naa15-bound) and uncomplexed Naa10 of *S. pombe*. The complexed form of Naa10 adopts a GNAT fold containing a central acetyl CoA–binding region and flanking N- and C-terminal segments that allows the acetylation of conventional substrates (Liszczak et al., 2013). In the uncomplexed form, Leu22 and Tyr26 shift out of the active site of Naa10, and Glu24 is repositioned by ∼5 Å resulting in a conformation that presumably allows for the acetylation of acidic N-termini (Liszczak et al., 2013). However, some proteins starting with an N-terminal acidic amino acid are usually further modified by arginyl-transferases and targeted by the Arg/N-end rule pathway for degradation (Varshavsky, 2011). Therefore, further studies have to show if Type I actins are unique substrates of non-complexed Naa10 and/or if more *in vivo* substrates with acidic N-termini exist. Such studies also need to explore whether the NTA of actin does trigger any downstream processing in the Arg/N-end rule pathway.

Apart from its function as an N-terminal acetyltransferase, NatA has been shown at least *in vitro* to possess N-terminal propionyltransferase activity (Foyn et al., 2013b) and lysine acetylation activity (Jeong et al., 2002; Lin et al., 2004; Lim et al., 2006; Yoo et al., 2006; Lim et al., 2008; Lee et al., 2010b; Shin et al., 2014). Autoacetylation at an internal lysine K136 in human and mouse Naa10 has been shown to regulate its enzymatic activity (Seo et al., 2010; Seo et al., 2014). Particularly, in *in vitro* acetylation assays with recombinantly expressed and purified wt Naa10 and subsequent detection with anti-acetyl-lysine antibody revealed acetylation of mNaa10^225^, mNaa10^235^ and hNaa10^235^ (see below for isoforms), whereas Naa10 with K82A and/or Y122F mutations, that inhibit acetyl-CoA binding, or Naa10 K136 (a mutation of the putative acetyl-acceptor site) were not found to be acetylated (Seo et al., 2014). In strong contrast to this, LC/MS/MS analyses on human Naa10 expressed and purified from *E. coli* did not show acetylation at any of the internal 16 lysines after incubation with Acetyl-CoA (Murray-Rust et al., 2006). Generally, ε-acetylation of internal lysines by Naa10 and other NATs is quite controversial in the field, and structural comparisons suggests that specific loops (β6-β7 hairpin loops in NatA and NatE or an extended α1-α2 loop in NatD) which are absent in KATs (lysine acetyltransferase) would block internal lysines from being inserted into the active site (Magin et al., 2015). Thus it is not yet known how much lysine is directly acetylated by Naa10 *in vivo*, and the degree to which autoacetylation of Naa10 occurs *in vivo* is also currently not well characterized.

#### 1.2.1 Mammalian Naa10 and isoforms

Naa10 (N^α^-acetyltransferase 10; NatA catalytic subunit; ARD1, arrest-defective protein 1 homolog; DXS707; TE2), the catalytic subunit of NatA, has an apparent molecular weight of 26 kDa and contains a typical Gcn5-related N-acetyltransferases (GNAT) domain. In mouse, *NAA10* is located on chromosome X A7.3 and contains 9 exons. Two alternative splicing products of mouse Naa10, mNaa10^235^ and mNaa10^225^, were reported in NIH-3T3 and JB6 cells that may have different activities and function in different subcellular compartments (Chun et al., 2007). The human *NAA10* is located on chromosome Xq28 and is encoded by 8 exons (Tribioli et al., 1994). According to RefSeq (NCBI) (Pruitt et al., 2007), three different isoforms, Naa10^235^, Naa10^220^ and Naa10^229^, derived from alternate splicing (see Figure 2) exist. Recently, an additional putative splice variant, hNaa10^131^ (GenBank accession no. BC063377), was identified by Sanger sequencing in a single clone after amplification of Naa10 from HeLa cDNA (Seo et al., 2015). However, this isoform is identical to Naa10 isoform 2, except that the splice acceptor site is shifted by 1 bp, which results in a frameshift in the resulting mRNA and a premature translational stop. Since this variant is neither annotated at NCBI, UniProt or Ensembl and was not detected on the protein level, we will not further discuss implications of this putative variant. Additionally, a processed *NAA10* gene duplicate *NAA11* (ARD2) has been identified that is expressed in several human cell lines including Jurkat, HEK293 and NPA (Arnesen et al., 2006b). However, later studies have revealed data arguing that Naa11 is not expressed in the human cell lines HeLa and HEK293 or in cancerous tissues, and *NAA11* transcripts were only detected in testicular and placental tissues (Pang et al., 2011). Therefore, the functional role of Naa11 might be constricted to certain tissues only. Naa11 has also been found in mouse, where it is mainly expressed in the testis (Pang et al., 2009). *NAA11* is located on chromosome 4q21.21 or 5 E3 for human or mouse, respectively, and only contains two exons.

**Figure 2:**
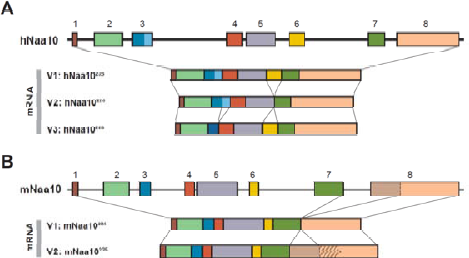
*NAA10* transcript variants. A) Human *NAA10* according to RefSeq. There are 3 human transcript variants. Variant 1 represents the longest transcript and encodes the longest isoform (235 aa). Variant 2 lacks in-frame exon 6 in the coding region and is shorter than isoform 1 (220 aa). Variant 3 uses an alternate in-frame splice site in exon 3 (229 aa). B) In mouse, two *NAA10* transcripts are described. Variant 1 represents the shorter transcript but the longer isoform (235 aa). Variant 2 uses an alternate splice site (exon 8), which results in a frameshift that induces a stop codon (*). The resulting isoform has a shorter and distinct C-terminus.

#### 1.2.2 Mammalian Naa15 and isoforms

Human Naa15 (N^α^-acetyltransferase 15; NatA auxiliary subunit, NMDA Receptor-Regulated Protein; NARG1; Tubedown-100; Tbdn100; tubedown-1, NATH) is well described as the auxiliary subunit of NatA, although it may have NatA-independent functions. The human *NAA15* gene is located on chromosome 4q31.1 and contains 23 exons. Initially, 2 mRNA species were identified, 4.6 and 5.8 kb, both harboring the same open reading frame encoding a putative protein of 866 amino acids (∼105 kDa) protein that can be detected in most human adult tissues (Fluge et al., 2002). According to RefSeq/NCBI (Pruitt et al., 2007), only one human transcript variant exists, although 2 more isoforms are predicted. In addition to full length Naa15, a N-terminally truncated variant of Naa15 (named tubedown-1), Naa15_273-865,_ has been described (Gendron et al., 2000). However, northern blot analyses of poly(A) mRNA from mouse revealed that full length Naa15 is widely expressed, whereas smaller transcripts were visualized only in heart and testis (Willis et al., 2002).

Similar to the situation with *NAA10*, a *NAA15* gene variant has been identified, *NAA16*, that originates from an early vertebrate duplication event (Arnesen et al., 2009a). The encoded protein shares 70% sequence identity to hNaa15 and is expressed in a variety of human cell lines, but is generally less abundant as compared to hNaa15 (Arnesen et al., 2009a). Three isoforms of Naa16 are validated so far (NCBI RefSeq). Mouse *NAA15* is located on chromosome 2 D and contains 20 exons, whereas mouse *NAA16* is located on chromosome 14 D3 and consists of 21 exons.

It has been shown in principle that NatA could assemble from all the isoforms. Naa15 interacts with Naa11, in humans (Arnesen et al., 2006b) and mouse (Pang et al., 2009), and Naa10 interacts with the Naa15 paralogue, Naa16, creating a more complex and flexible system for N^α^-terminal acetylation as compared to lower eukaryotes (Arnesen et al., 2009a). Such a system might create the opportunity for functional redundancy or compensation in the event of loss of Naa10 or Naa15, although we are not aware of any studies showing whether *NAA11* expression might be upregulated in tissues lacking or having reduced *NAA10* expression.

#### 1.2.3 Other NatA components

Many studies document the interaction between Naa10 and Naa15 and since a crystal structure of *S. pombe* Naa10 bound to Naa15 has been published recently, it is safe to conclude that this is a very stable complex. We await such crystal structures for human Naa10 and Naa15. As an enzyme, NatA transiently interacts with a variety of substrates *per se*. However, some evidence indicates that NatA constitutively interacts with specific proteins to assemble into trimeric or even larger complexes, which might modulate NatA function. One such example is Naa50 (N^α^-acetyltransferase 50; NatE; NAT13; Mak3; Nat5, SAN). Naa50 is the catalytic acetyltransferase subunit of NatE, is expressed in several human cell lines, and has been shown to be associated with NatA in yeast (Gautschi et al., 2003), fruit fly (Williams et al., 2003) and humans (Arnesen et al., 2006a). Naa50 has a distinct substrate activity for Met followed by a hydrophobic amino acid in human and yeast (Polevoda et al., 1999; Evjenth et al., 2009; Evjenth et al., 2012) and has been claimed to possess ε-acetyltransferase activity towards K525 in β-tubulin (Chu et al., 2011) and histone 4 (Evjenth et al., 2009). Furthermore, hNaa50 has been shown to harbor autoacetylation activity on internal lysines (K34, K37 and K140) *in vitro*, modulating Naa50 substrate activity (Evjenth et al., 2009; Evjenth et al., 2012). However, a recent structural study on human Naa50 contradicts these findings. The X-ray crystal structure of human Naa50 revealed, similar to Naa10, a GNAT fold with a specific substrate binding groove that allows acetylation of α-amino substrates but excludes lysine side chains (Liszczak et al., 2011). This seems to be strong evidence against a role for Naa50 in direct acetylation of lysine side chains. Further studies have to sort out these discrepancies.

Because Naa50 has a distinct/different substrate specificity and *NAA50*Δ cells did not display the NatA phenotype in yeast (Gautschi et al., 2003), Naa50 was considered as an independent NAT and was named NatE (Starheim et al., 2009b). Furthermore, in HeLa cells, more than 80 % of endogenous Naa50 is not associated with the NatA complex (Hou et al., 2007). Therefore, future experiments have to examine whether Naa50 has a distinct function independent of NatA and/or if Naa50 works in a cooperative manner with NatA.

Recently, the chaperone like protein HYPK (Huntingtin Interacting Protein K) was shown to interact with Naa10 and 15 and is required for NTA of the known *in vivo* NatA substrate PCNP (Arnesen et al., 2010). However, it is an open question whether HYPK generally is required for NatA-mediated acetylation of downstream substrates.

Further interaction partners of Naa10 have been identified including Myosin light-chain kinase (Shin et al., 2009), tuberous sclerosis 2 (Kuo et al., 2010) RelA/p65 (Xu et al., 2012) DNA methyltransferase 1 (Lee et al., 2010a), androgen receptor (Wang et al., 2012) and proteasome activator 28β (Min et al., 2013). In high throughput screens cell division cycle 25 homolog (Rual et al., 2005) and Rho guanine nucleotide exchange factor 6 have been shown as interaction partners of NatA (Xiao et al., 2007). Additionally, β-Catenin (Lim et al., 2006; Lim et al., 2008), HIF-1α (Jeong et al., 2002; Arnesen et al., 2005b) and methionine sulfoxide reductase A (Shin et al., 2014) have been suggested to bind to Naa10. In a recent high-throughput study, multiple orthogonal separation techniques were employed to resolve distinct protein complexes. Fractionation of soluble cytoplasmic and nuclear extracts from HeLa S3 and HEK293 cells into 1,163 different fractions identified several interaction partners for Naa10 (Naa15, Naa16, Mina, M89BB, TCEA1 and PLCβ3), Naa15 (RT21, ML12A, HYPK and Cap1) and Naa16 (TCEA1, PLCB3, Naa10 and Mina) (Havugimana et al., 2012). However, these interactions seem to be transient in the cell and have not yet been shown to regulate or change NatA function; therefore, we do not list them as part of any putative larger NatA complex. Evidence of a larger stable complex, other than the dimer between Naa10 and Naa15, could come from structural studies, including possibly cryo-electron microscopy.

#### 1.2.4 Localization of NatA

Mainly from yeast data, it is thought that the auxiliary subunits of NatA as well as other NATs are associated with mono- and polysome fractions and co-translationally acetylate the nascent polypeptide chain as it emerges from the ribosome (Gautschi et al., 2003; Polevoda et al., 2008). In line with this, it has been shown that human Naa10 and Naa15, HYPK (Arnesen et al., 2010), the human paralog of Naa15, Naa16 (Arnesen et al., 2009a) as well as yeast Naa15 (Raue et al., 2007) and rat Naa15 (Yamada and Bradshaw, 1991) are associated with poly- or monosomes. In yeast, NatA binds via the ribosomal proteins, uL23 and uL29 (Polevoda et al., 2008). Further data indicates that NatA preferably associates with translating ribosomes. Particularly, yNatA as well as other ribosome-associated protein biogenesis factors (including the chaperones Ssb1/2 and ribosome-associated complex, signal recognition particle and the aminopeptidases Map1 and Map2) bind with increased apparent affinity to randomly translating ribosomes as compared with non-translating ones (Raue et al., 2007). Hsp70 chaperones may be direct targets of NatA, and NTA by NatA contributes an unanticipated influence on protein biogenesis, both through and independent of Hsp70 activity (Holmes et al., 2014), supporting a role of NatA in protein biogenesis. However, the NatA complex also exists in a ribosome-free context. For instance it has been shown that the majority of hNatA is non-polysomal (Arnesen et al., 2005a) and a minor fraction of cytosolic hNaa10 exists independent of the NatA complex, carrying out post-translational acetylation as mentioned above (Van Damme et al., 2011b). Mammalian Naa10, Naa11 and Naa15 and Naa50 (isoforms) have been reported to be mainly localized in the cytoplasm and to a lesser extent to the nucleus (Fluge et al., 2002; Sugiura et al., 2003; Bilton et al., 2005; Arnesen et al., 2006a; Arnesen et al., 2006b; Chun et al., 2007; Xu et al., 2012; Park et al., 2014; Zeng et al., 2014; Aksnes et al., 2015c). In mouse, an isoform specific localization of Naa10 has been described. mNaa10^235^ was mainly nuclear in NIH-3T3 and JB6 cells whereas another variant mNaa10^225^, derived from alternative splicing at a different 3’-splice site, was mainly localized in the cytoplasm (Chun et al., 2007). In humans, Naa10^225^ is absent and Naa10^235^ was found to be evenly distributed in both cytoplasm and nucleus of HeLa and HT1080 cells as seen by immunofluorescence, confocal microscopy, and cell fractionation (Chun et al., 2007). Naa10 could be detected in nuclear fractions of doxorubicin treated HEK293 cells whereas a deletion construct lacking amino acids 1-35 could not be detected suggesting that a nuclear localization signal (NLS) resides in the N-terminal part of Naa10 (Park et al., 2012). Sequence analysis had previously identified a putative NLS more C-terminally in Naa10 between residues 78 and 83 (KRSHRR) (Arnesen et al., 2005a). In agreement with that, deletion of this NLS_78-83_ almost completely abrogated nuclear localization of Naa10, whereas Naa10 wild type was imported to the nuclei of proliferating HeLa and HEK293 cells, especially during S phase (Park et al., 2014). Furthermore, the deletion of NLS_78-83_ altered the cell cycle and the expression levels of cell cycle regulators and resulted in cell morphology changes and cellular growth impairment, all of which was mostly rescued when the nuclear import of hARD1 was restored by exogenous NLS (Park et al., 2014). Also, Arnesen *et al.* reported that neither leptomycin B nor actinomycin D significantly changed the localization patterns of Naa10 in HeLa cells, indicating that Naa10 is not actively imported through importin β-dependent mechanisms (Arnesen et al., 2005a).

In conclusion, it is well established that Naa10 localizes to the cytoplasm and the nucleus and the possible shuttling between the two compartments might be regulated by an internal NLS. Although the action of cytosolic Naa10 is well described in co-translational acetylation, the significance of its nuclear localization is still not well characterized; however, some studies connect Naa10 to nuclear processes such as transcriptional regulation (see below). Also, the signal pathways that might induce or regulate cytosolic to nuclear translocation are unknown.

Naa15 also harbors a putative NLS between residues 612-628 (KKNAEKEKQQRNQKKKK); however, only Naa10 was found to be localized in the nuclei of HeLa, GaMg, HEK293, MCF-7 and NB4 cells, whereas Naa15 was predominantly localized in the cytoplasm (Arnesen et al., 2005a). In contrast to this, a different study showed that Naa15 localizes to the nucleus where it interacts with the osteocalcin promoter, as shown by cellular fractionation and ChIP experiments in MC3T3E1 calvarial osteoblasts (Willis et al., 2002). Further studies have to resolve these discrepancies and analyze possible cell-type specific differences.

Besides that, it has been shown that Naa10 associates with microtubules in dendrites in cultured neurons (Ohkawa et al., 2008) and Naa15 colocalizes with the actin-binding protein cortactin and the F-actin cytoskeleton in the cytoplasm of IEM mouse and RF/6A rhesus endothelial cells (Paradis et al., 2008).

## 2 mammals

### 2.1 Naa10 function in mammals

The N^α^-acetyltransferase NatA is expressed widely in many tissues and NatA N-termini are overrepresented in eukaryotic proteomes. As pointed out earlier, 80-90% of soluble human proteins are fully or partially acetylated and nearly 40-50% of all proteins are potential NatA substrates according to their sequence in *S. cerevisiae*, *D. melanogaster* and humans (Van Damme et al., 2011c; Starheim et al., 2012). Due to its ubiquitous expression in almost all tissues and the broad substrate specificity of Naa10, it is perhaps not surprising that Naa10 might regulate protein function, stability and/or complex formation on a variety of substrates and subsequently modulate many cellular processes including cell cycle regulation, cancer progression, DNA damage, stress response, development and disease.

In the current model, Naa15 links Naa10, Naa50 and possibly other factors like HYPK to the ribosome where NatA/Naa10 acetylates the N^α^-amino group of canonical substrates and NatE/Naa50 acetylates methionine followed by a hydrophobic amino acid in a co-translational manner (Figure 3). Some known canonical NatA substrates include PCNP (Arnesen et al., 2010), androgen receptor (Wang et al., 2012), caspase-2 (Yi et al., 2011), α-tubulin (Ohkawa et al., 2008) and TSC2 (Kuo et al., 2010), although a wealth of proteomic studies in recent years have suggested many more substrates (Arnesen et al., 2009b; Van Damme et al., 2011a; Lange et al., 2014). Post-translational acetylation by non-ribosome-associated NatA or monomeric Naa10 might occur as well.

**Figure 3:**
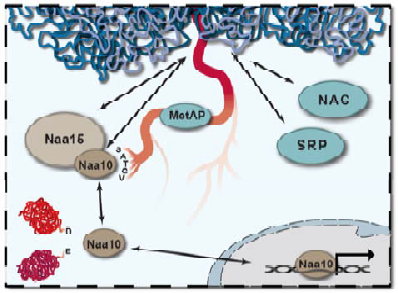
Multiple functions of Naa10. Associated with the ribosome in the NatA complex, Naa10 co-translationally acetylates the N^α^-terminal amino group of the nascent polypeptide chains of classical substrates as they emerge from the ribosome. Naa15 as well as the signal recognition particle SRP and nascent polypeptide-associated complex NAC might bind competitively to similar regions on the ribosome near the exit tunnel (see below). This might indicate a general involvement of Naa10 in protein biogenesis. Uncomplexed Naa10 post-translationally N^α^-acetylates proteins starting with acidic side chains and might also N^ε^-acetylate internal lysines. Furthermore, it has been suggested that Naa10 translocates into the nucleus where it acts in cooperation with transcription factors to modulate protein expression.

On the other hand, when Naa10 is not in a complex with Naa15, Naa10 adopts a different catalytic activity towards N-termini with acidic side chains. Presumably, this function occurs post-translationally in the cytosol, as Naa15 is necessary to link Naa10 to the ribosome (Figure 3). To date, only two substrates with acidic N termini, γ- and β-actin, have been described (Van Damme et al., 2011b). Proteins that have been reported to be substrates for ε-acetylation at lysines by Naa10 include β-catenin (Lim et al., 2006; Lim et al., 2008), Runx2 (Yoon et al., 2014), MSRA (Shin et al., 2014), MLCK (Shin et al., 2009), androgen receptor (Wang et al., 2012) or HIF-1α (Jeong et al., 2002; Yoo et al., 2006; Lee et al., 2010b) although there is controversy in the field whether HIF-1α is an actual substrate of Naa10 or not (see below). Another ambiguous finding is that Naa10 can translocate into the nucleus, regulating gene transcription, and this function might even be independent of its catalytic activity (Figure 3). For example, in H1299 lung cancer cells, Naa10 binds to nonmethylated DNA at the E-Cadherin promoter, directly interacts with DNMT1 (DNA methyltransferase 1) thereby recruiting DNMT1 resulting in silencing of the E-Cadherin promoter in a NAT-independent manner as analyzed with a Naa10-R82A mutant (Lee et al., 2010a). Similarly, the enzymatic activity is not necessary for Naa10 to significantly suppress migration, tumor growth, and metastasis in human cancer cells. Instead, Naa10 binds to the GIT-binding domain of PIX (Rho guanine nucleotide exchange factor 7), thereby preventing the formation of the GIT-PIX-Paxillin complex, resulting in reduced intrinsic Cdc42/Rac1 activity and decreased cell migration (Hua et al., 2011).

As indicated above, Naa10 has been implicated in the regulation of gene transcription and cell motility. In this regard, it has been shown that overexpression or knock-down of Naa10 in HT1080 cells reduces migration and enhances invasion (Shin et al., 2009). The authors also reported that Naa10 interacts with activated (phosphorylated) MYLK (myosin light-chain kinase) and acetylates it at Lys^608^, thereby inactivating MYLK resulting in the dephosphorylation of MLC (Shin et al., 2009). Additionally, many other functions of Naa10 have been discussed including cyclin D1 regulation, regulation of DNA-damage response pathways, cellular hypoxia, apoptosis and cancer.

In *S. cerevisiae,* NatA function is not essential but disruption has other strong defects (see below). However, the *D. melanogaster* homolog of Naa10 (variable nurse cells; vnc) was found to be crucial for normal development and loss of Naa10 results in lethality. Particularly, disruptions of this gene result in pleiotropic oogenesis defects including abnormal cyst encapsulation, desynchronized cystocyte division, disrupted nurse cell chromosome dispersion, and eventual lethality for the animal, with homozygotes for *vncBDk (NAA10*Δ) perishing during the second larval instar (Wang et al., 2010). Additionally, Naa10 is essential in controlling *C. elegans* life history (see below) and loss of the corresponding homologs in *T. brucei* is lethal as well (Ingram et al., 2000; Chen et al., 2014). Surprisingly, Naa10-knockout mice have very recently been reported to be viable, displaying a defect in bone development (Yoon et al., 2014). However, a full characterization of the Naa10 knockout in mouse remains to be published. Below, we will summarize and discuss the recent findings on Naa10 function. For a recent review on Naa10 function in cancer we refer the reader to (Kalvik and Arnesen, 2013).

#### 2.1.1 Cyclin D1 regulation

Recent studies link Naa10 activity to several signaling pathways, including Wnt/β-catenin, MAPK and JAK/STAT signaling, that regulate cyclin D1 expression and cell cycle control. The proto-oncogene cyclin D1 forms a complex with CDK4/6 (cyclin-dependent kinase 4/6) and promotes G1/S cell cycle transition. The expression of cyclin D1 is tightly regulated by many factors including growth factor-dependent activation of the canonical MAPK (mitogen-activated protein kinase) pathway that stimulate the expression of AP-1 transcription factors (c-Fos and c-Jun), NF-κB signaling, cytokines through the JAK-STAT pathway and Wnt-signaling (Klein and Assoian, 2008).

The Wnt signal transduction pathway is evolutionarily conserved and regulates many cellular functions including cell migration, cell polarity and embryonic development. In the canonical Wnt pathway, cytoplasmic β-catenin is marked for ubiquitination and proteasomal degradation by the β-catenin destruction complex composed of axin, APC (adenomatous polyposis coli), PP2A (protein phosphatase 2A), GSK3 (glycogen synthase kinase 3) and CK1α (casein kinase 1α) in the absence of stimuli (Niehrs, 2012). Activation of Frizzled and its coreceptor LRP5/6 by secreted glycoproteins called Wnt disrupts the destruction complex, leading to a stabilization and accumulation of β-catenin that translocates to the nucleus where it participates in the regulation of LEF/TCF (Lymphoid enhancer-binding factor 1/T-cell factor) dependent genes like c-Myc, cyclin D1 or c-jun (Komiya and Habas, 2008). c-Myc inhibits the transcription of the cyclin kinase inhibitor p21^(WAF1/CIP1)^ thereby activating cyclin-dependent kinases and promoting cell cycle progression (Gartel et al., 2001).

The first evidence for an implication of Naa10 in the regulation of cyclin D1 came from studies on the small lung cancer cell lines H1299 and A549. Knockdown of Naa10 by siRNA in these cells attenuated β-catenin acetylation, reduced the recruitment of β-catenin/TCF (T-cell factor) to the cyclin D1 promoter in chromatin immunoprecipitations, and diminished the transactivation activity of a TCF-reporter assay (TOP-FLASH) resulting in a reduced cyclin D1 expression and proliferation inhibition (Lim et al., 2006). The direct interaction of Naa10 and β-catenin and the ε-acetylation of β-catenin at lysine(s) could be shown in pulldown and *in vitro* acetylation assays with ectopically expressed proteins, respectively (Lim et al., 2006). A follow up study from the same group showed that this regulation has implications in hypoxic stress management. Naa10 forms a stable complex with β-catenin and acetylates β-catenin, which leads to the activation of TCF4 dependent genes, promoting cell proliferation under normoxic conditions, whereas HIF-1α dissociates the β-catenin/Naa10 complex in hypoxia (Lim et al., 2008). Particularly, HIF-1α competes via its oxygen-dependent degradation domain (ODDD) with β-catenin for Naa10 binding, thus leading to hypoacetylation and a repressed transcriptional activity of β-catenin including p21^WAF1/CIP1^-dependent growth arrest (Lim et al., 2008). In contrast to this, no changes in β-catenin expression or acetylation could be observed upon siRNA mediated depletion of Naa10 in CAL62 and 8305C cells (Gromyko et al., 2010). CAL-62 and 8305C cells, bearing p53 mutations, show little intrinsic β-catenin/TCF activity (Adam et al., 2012), whereas H1299 and A549 cells have activated β-catenin (Lim et al., 2006; Lim et al., 2008). This could indicate that activation of the β-catenin/TCF pathway is an essential requirement or prerequisite for Naa10 to acetylate β-catenin. However, another mechanism by which HIF could block β-catenin signaling without affecting β-catenin acetylation status has been described: HIF-1α N-terminal domain directly competes with TCF4 for β-catenin complex formation, thereby inhibiting β-catenin/TCF4 transcriptional activity, resulting in a G1 arrest that involves c-Myc and p21^WAF1/CIP1^ under hypoxic conditions (Kaidi et al., 2007). Therefore, it has been speculated that the hypoxic repression of TCF4 is subject to double-checking by (at least) two distinct mechanisms: competitive disruption of the Naa10/β-catenin complex and disruption of the β-catenin/TCF4 transcriptional complex (Lim et al., 2008).

The enzymatic activity of Naa10 seems to be important for its ability to stimulate β-catenin/TCF transcriptional activity. Seo and colleagues showed that Naa10 undergoes autoacetylation at an internal lysine K136, which in turn increases its enzymatic activity towards other substrates, including β-catenin (Seo et al., 2010). Naa10 wild type augments LiCl-induced recruitment of β-catenin to the cyclin D1 promoter, amplifies TCF4 reporter in HEK293 cells, enhances cell growth/increased cell proliferation in A549, HeLa, and HEK293T cells, thereby leading to increased colony formation in an anchor-independent colony formation assay and increased tumor growth in mouse xenografts with H460 human lung cancer cells expressing wild type Naa10 (Seo et al., 2010). An autoacetylation-deficient mutant K136R mutant abrogated these effects indicating that the enzymatic activity of Naa10 and the resulting auto-acetylation is important for its signaling (Seo et al., 2010). In line with this, knockdown of endogenous Naa10 decreased luciferase activity in a cyclin D1 reporter system and reduced cyclin D1 expression, which could be restored by Naa10 wild type but not by the K136R mutant (Seo et al., 2010). In mouse, an isoform specific regulation has been described. Overexpression of mNaa10^235^ increased Cyclin D1 levels as shown by RT-PCR and western blot whereas overexpression of wt mNaa10^225^ or mutated mNaa10^235^ that abrogate acetyl-CoA binding or autoacetylation (K82A/Y122F or K136R) had no effects (Seo et al., 2014).

As mentioned above, Naa10-depletion has been shown to decrease cyclin D1 levels and inhibit cell cycle promotion independently of β-catenin in CAL62 and 8305C cells (Gromyko et al., 2010). The authors speculate that Naa10 knockdown might instead induce the DNA-damage response network and p53-dependent apoptosis independent of β-catenin (Gromyko et al., 2010). In this regard, it should be mentioned that Naa10 might regulate cyclin D1 expression also via the MAPK (mitogen-activated protein kinase)-pathway. Phosphorylation/activation of the MAPK pathway activates Erk1/2 (extracellular signal-regulated kinase 1/2) and stabilizes c-Fos by direct acetylation, which then associates with c-Jun to form the transcriptionally active AP-1 (activator protein) complex that activates cyclin D1 (Cargnello and Roux, 2011). Knockdown of Naa10 significantly decreases phorbol ester TPA-induced phosphorylation of Erk1/2, attenuates c-Jun and c-Fos activation, decreases AP-1-recruitment to the cyclin D1 promoter and results in a repression of the AP-1 target genes including cyclin D1 (Seo et al., 2010). Also, Naa10 has been shown to interact with STAT5a in *in vitro* association and immunoprecipitation assays, thereby inhibiting NF-κB-dependent *IL1B* (interleukin-1β) expression and reducing STAT5a-dependent *ID1 (*inhibitors of differentiation 1) expression (Zeng et al., 2014), which in turn could regulate cyclin D1 expression, as cytokines, such as interleukin-3 and interleukin-6, stimulate cyclin D1 promoter activity via STAT3 and STAT5 (Klein and Assoian, 2008). However, this has not been tested in the above study (Zeng et al., 2014).

Aside from cyclin D1, Naa10 has been linked to other factors implicated in cell-cycle and cell proliferation regulation. Naa10 has, for example, been identified in a yeast two-hybrid screen as an interaction partner of CDC25A (cell division cycle 25 homolog) that is required for G1/S cell cycle progression (Rual et al., 2005). Furthermore, Naa10 has been shown to physically interact with TSC2 (tuberous sclerosis 1-2 complex), an inhibitor of mTOR (mammalian target of rapamycin) (Kuo et al., 2010). Particularly, Naa10-dependent acetylation of TSC2 induced the stabilization of TSC2, repression of mTOR activity leading to reduced cell proliferation and increased autophagy of MCF-10A and MDA-MB-435 cells (Kuo et al., 2010). However, it should also be noted that other studies postulate that Naa10 does not regulate cell cycle progression, e.g. deficiency of Naa10/Naa15 had no effect on cell-cycle progression in HeLa and U2OS cells (Yi et al., 2011).

To summarize, Naa10 seems to activate and/or amplify the transcriptional activity of β-catenin/TCF transcriptional activity thereby stimulating cyclin D1 and c-Myc expression leading to inhibition of p21^WAF1/CIP1^ and promoting the G1/S cell cycle transition. The mechanism of such action is not clear to date and there are some discrepancies whether Naa10 acetylates β-catenin directly; however, the catalytic activity of Naa10 seems to be important for this function. However, accumulating evidence suggests that Naa10 only exerts effects when the Wnt/β-catenin pathway is either intrinsically or extrinsically activated, e.g. though stimulation with LiCl. In line with that, Naa10 knockout did not affect β-catenin ubiquitination or degradation (Lim et al., 2006). This indicates that Naa10 is rather a positive regulator than an activator of the Wnt/β-catenin pathway and that the cellular context determines the action of Naa10. Similarly, other cellular conditions such as hypoxia might influence regulatory functions of Naa10 and some data suggests that Naa10 regulates cyclin D1 through other pathways, including MAPK and JAK/STAT. Future studies have to examine the acetylation status and the consequences of this modification on the involved proteins under various cellular conditions in detail.

#### 2.1.2 NF-kB/ DNA damage

Data generated from yeast indicates that NatA is involved in chromosomal stability, telomeric silencing and NHEJ repair (Aparicio et al., 1991; Ouspenski et al., 1999; Wilson, 2002). Indeed, a genome-wide RNA interference screen in *D. melanogaster* cells identified Naa10 as a regulator of apoptosis as a response to doxorubicin-induced DNA damage (Yi et al., 2007). Furthermore, Naa10 regulates cell death in response to doxorubicin in HeLa, HT1080, and U2OS cells (Yi et al., 2011) and apoptosis in response to the DNA damage inducer 5-fluorouracil in RKO and H1299 cells (Xu et al., 2012), indicating a function of Naa10 in DNA damage control.

The dimeric transcription factor NF-κB (nuclear factor κB) is activated by DNA-damaging drugs such as doxorubicin, daunorubicin and mitoxantrone, that intercalate into DNA and inhibit topoisomerase II as part of the cellular stress response (Karl et al., 2009). To date, five members of the NF-κB family are known: RelA (p56), RelB, c-Rel, NF-κB1 (p105/p50) and NF-κB2 (p100/52). In unstimulated cells, the IκB (Inhibitor of κB) proteins, mask the nuclear localization signals of NF-κB proteins, thereby keeping them sequestered in an inactive complex in the cytoplasm (Huxford et al., 1998; Jacobs and Harrison, 1998). Activation of the classic/canonical NF-κB pathway by cytokines such as IL-1 (interleukin-1) or TNFα (tumor necrosis factor α) leads to the recruitment of several factors, including RIP1 (receptor-interacting protein 1), TRAF2 (TNF receptor-associated factor 2) and cIAPs (inhibitor of apoptosis proteins) that results in the activation of IKK (IκB kinase), a complex of IKKα, IKKβ and NEMO (NFκB essential modifier). This complex phosphorylates and marks IκB for ubiquitination and subsequent proteasomal degradation, which in turn releases and activates NF-κB that then translocates into the nucleus and induces the transcription of downstream target genes (Hayden and Ghosh, 2012). In DNA damage, RIP1 forms a distinct complex with PIDD (P53 inducible death domain-containing protein) and NEMO in the nucleus. NEMO is subsequently SUMOylated and phosphorylated, resulting in the nuclear export of NEMO where it forms the IKK complex and activates NF-κB (McCool and Miyamoto, 2012).

Park and colleagues showed that siRNA-mediated knockdown attenuated, or overexpression of Naa10 increased, respectively, doxorubicin-induced RIP1/PIDD/NEMO complex formation, NEMO ubiquitination and NF-κB activation in HEK293 cells (Park et al., 2012). Specifically, Naa10 binds RIP1 via its acetyltransferase domain (aa45–130); however, the N-terminus as well a functional active acetyltransferase domain of Naa10 are necessary to induce NF-κB activation (Park et al., 2012). TNFα-induced NFκB activation was not affected by Naa10 knockdown (Yi et al., 2011; Park et al., 2012). In line with that, *in vitro* pull-down and co-immunoprecipitation assays in three cancer cell lines (RKO, H1299 and A549) showed that Naa10 directly interacts with NF-κB (RelA/p56) and contributes in NF-κB transcriptional activation of MCL1 (induced myeloid leukemia cell differentiation protein 1), thereby protecting cells from apoptosis (Xu et al., 2012). In contrast to this, a very recent report showed that overexpression or siRNA-mediated knockdown of Naa10 had no effects on cisplatin-induced DNA damage in A549 and H1299 cells (Shin et al., 2014), suggesting that Naa10 function in DNA damage depends on the stimulus as well as the cellular system.

On the other hand, Naa10 might also induce caspase-dependent cell death upon treatment with DNA-damaging drugs. In this regard it has been shown that Naa10 is essential for the activation of caspase-2/-3/-7 and -9 in HeLa cells after doxorubicin stimulation (Yi et al., 2007). Later studies by the same group showed that Naa10 directly acetylates Caspase-2, which induces its interaction with caspase-2 scaffolding complex RAIDD (RIP-associated ICH-1/CED-3 homologous protein with a death domain), and activation of Caspase 2 (Yi et al., 2011) thereby inducing apoptosis in response to DNA damage. Another study showed that Naa10 associates with IκB kinase β (IKKβ) which phosphorylated Naa10 at Ser209, resulting in destabilization and proteasome-mediated degradation of Naa10 (Kuo et al., 2009), supporting the direct link between Naa10 and the NF-κB pathway and providing a possible feedback regulation of Naa10 during NF-κB signaling. In line with that, a recent study found that Runx2, a transcription factor in bone development, stabilizes Naa10 by inhibiting IKKβ-dependent phosphorylation and degradation of Naa10 (Yoon et al., 2014). Naa10 itself is degraded only at late stages of DNAdamage-induced apoptosis in HeLa and HIH-3T3cells (Chun et al., 2007), which also might be a hint that it is important at early stages of DNA damage regulation.

Naa10 might also directly induce DNA-damage response pathways. hNatA-depletion in HCT116 cells, for instance, itself activates DNA damage signaling through the γH2AX and Chk2 sensing system, leading to p53 dependent apoptosis (Gromyko et al., 2010). In contrast to this, another study showed that Naa10 overexpression, but not siRNA-mediated knockdown, increased γH2AX expression in A549 and H1299 cells upon treatment with oxidative stress, but had no effect in untreated cells (Shin et al., 2014). The reason for these discrepancies is not clear.

However, siRNA mediated knockdown of Naa10 in HeLa cells leads to apoptosis and sensitizes cells for daunorubicin-induced apoptosis (Arnesen et al., 2006c) and knockdown of Naa10 in the cancer cell line H1299 cells augmented the cellular response to doxorubicin (cell death and caspase-3 activation) (Lim et al., 2008). This is in clear contrast to before, as knockdown of Naa10 dramatically enhanced cell survival in the presence of doxorubicin in HeLa and *D. melanogaster* Kc_167_ cells (Yi et al., 2007). These observed differences might be due to the context dependent regulation of NF-κB signaling and be dependent on the dose and duration of the applied drug. Recent reports show that, while DNA damage dependent NF-κB activation mediates protection of normal and malignant cells from DNA damage-induced apoptotic death, prolonged treatment or high dosing of cells with genotoxic chemotherapeutics rather induces apoptosis (involving ubiquitination of RIP1 and assembly of RIP1/NEMO/FADD/caspase-8) (McCool and Miyamoto, 2012).

However, Naa10 intrinsically is essential for cell survival in higher eukaryotes. Loss-of-function of Naa10 in Drosophila affects cell survival/proliferation and is lethal for the animal (Wang et al., 2010). Furthermore, knockdown of Naa10 impaired proliferation, induces cycle arrest and/or induces cell death as shown for different thyroid carcinoma and thyroid follicular epithelial cell lines (Gromyko et al., 2010), H1299 lung cancer cells (Lee et al., 2010a), LNCaP cells (Wang et al., 2012), HeLa (Arnesen et al., 2010) and RKO and H1299 cells (Xu et al., 2012). For a recent review on Naa10 in tumor progression see: (Kalvik and Arnesen, 2013)

In conclusion, despite efforts over the past 10 years, the role of Naa10 in DNA damage control pathways and NF-κB regulation is quite obscure, which is partly due to the fact that NF-κB regulation in response to DNA damage itself is not well understood. Different stimuli and even different concentrations of stimulus can have opposing effects downstream of NF-κB activation also depending on the cell system, ranging from protection of damaged cells from apoptotic cell death and DNA repair to cell cycle arrest, activation of caspase-8 and apoptosis. Similarly, Naa10-dependent effects in response to genotoxic stress vary strongly in the different studies. Naa10 seems to always enhance the effects with regard to NF-κB, whether it is induction of cell death or protection from apoptosis. More studies are necessary to draw exact conclusion on the mechanism, however, evidence suggests that Naa10 directly interacts with NF-κB and that a functional active acetyltransferase domain of Naa10 is necessary to induce NF-κB activation. Furthermore, two studies showed that phosphorylation of Naa10 by IKKβ induced proteasome-mediated degradation of Naa10 itself, which could indicate a possible feedback regulation for Naa10 in DNA-damage. However, again the significance of such an action is unclear. Aside from NF-κB, other pathways involved in DNA damage response, including γH2AX and the Chk2 sensing system, seem to be modulated by Naa10 leading to p53 dependent apoptosis. Furthermore, Naa10 has been shown to directly acetylate Caspase-2, directly implicating Naa10 in the activation of apoptosis. This clearly shows the complexity of Naa10 function in various pathways and highlights the importance of NTA research.

#### 2.1.3 Cellular hypoxia

The hypoxia-inducible factor (HIF) mediates the cellular response to low oxygen levels. HIF (hypoxia-inducible factors) is a heterodimeric transcription factor that regulates many cellular processes including proliferation, migration, differentiation, glucose metabolism and angiogenesis. HIF is composed of a constitutively expressed HIF-1β subunit and one of the three HIFα subunits (HIF-1α, HIF-2α, HIF-3α). Under normoxic conditions, HIFα is an exceptionally short-lived protein due to the O_2_-dependent hydroxylation of proline (Pro) or asparagine (Asn) residues by PHD (prolyl hydroxylase domain) proteins (Benizri et al., 2008). Particularly, hydroxylation of two conserved prolines (Pro_402_ and Pro_564_) in the oxygen-dependent degradation domain (ODDD) of HIF-1α recruits the E3-ubiquitin ligase, pVHL (von Hippel-Lindau), leading to ubiquitination and proteasomal degradation of HIF-1α. Additionally, hydroxylation of HIF-1α at Asn_903_ within the C-terminal transactivation domain (C-TAD) by another hydroxygenase termed FIH (factor inhibiting HIF) abrogates the binding of the essential transcriptional co-activators p300/CBP, and hence the transcriptional activity of the HIF-1α (Brahimi-Horn et al., 2007).

Under oxygen-limiting conditions, HIF-1α is stabilized, dimerizes with HIF-1β and binds to site-specific sequences termed hypoxia response elements (HRE), recruits the co-activator p300/CBP and regulates the transcription of specific target genes like VEGF, glycolytic enzymes and erythropoietin. Another regulation of HIF-1α that is independent of oxygen seems to involve Lysine K_532_ in HIF-1α (Tanimoto et al., 2000; Chun et al., 2003) and acetylation at this side chain accelerates HIF-1α proteasomal degradation (Demidenko et al., 2005).

Early studies showed that mouse Naa10 expression is induced by hypoxia and enhances the interaction of HIF-1α with VHL leading to HIF-1α ubiquitination and proteasomal degradation (Jeong et al., 2002). HIF-1α could be detected in immunoprecipitates with an acetyl-lysine antibody after induction of hypoxia or Naa10 transfection in HEK293 (Jeong et al., 2002), B16F10, MKN74 (Lee et al., 2010b) and MCF-7 (Yoo et al., 2006) cells, suggesting an *in vivo* acetylation of HIF-1α by Naa10, although this result was alternatively interpreted as an acetylated protein like p300/CBP or Hsp90 coprecipitating with HIF-1α (Fath et al., 2006). Nevertheless, it could be shown that Naa10 directly binds to the oxygen degradation domain (ODD) of HIF-1α and ε-acetylates HIF-1α at Lys532 *in vitro* (Jeong et al., 2002). Additionally, it has been shown that nickel (II) or cobalt (II) ions that are known to induce hypoxia-like stress, down-regulated Naa10 and FIH-1 mRNA in the human lung adenocarcinoma A549 cells (Ke et al., 2005). In contrast to this, several studies showed that human Naa10 expression is not affected in hypoxia and/or Naa10 does not acetylate and/or destabilize HIF-1α (Arnesen et al., 2005b; Bilton et al., 2005; Fisher et al., 2005; Fath et al., 2006; Murray-Rust et al., 2006). Particularly, neither mRNA nor protein levels of Naa10 are regulated by hypoxia in HeLa, HT1080, HEK293 or MCF-7 cells and overexpression or silencing of Naa10 has no impact on HIF-1α stability (Bilton et al., 2005). Fisher and colleagues also found no effect of Naa10 knockdown in HIF-1α protein level in HEK293T cells and HepG2 cells but detected that Naa10 is downregulated in a number of cell lines in response to hypoxia or CoCl_2_, a hypoxia mimic compound (Fisher et al., 2005). Furthermore, knockdown of Naa10 decreases erythropoietin and VEGF protein production under normoxic and hypoxic conditions and suppressed proliferation (Fisher et al., 2005). In contrast, Arnesen and colleagues did not find a decreased Naa10 protein level in hypoxia in HT1080, RCC4, HeLa, MCF-7 and HEK293 cells, but could confirm the interaction of Naa10 with HIF-1α ODD suggesting a putative, still unclear, connection between these proteins (Arnesen et al., 2005b).

More recently, studies suggested that different splicing forms from mouse and humans differentially regulate the cellular response to hypoxia. As mentioned earlier, 2 mouse (mNaa10^225^ and mNaa10^235^) and 3 human (hNaa10^235^, hNaa10^220^ and hNaa10^229^) isoforms of Naa10 that are generated by alternate splicing are listed in RefSeq (Pruitt et al., 2007). The used isoforms implicated in hypoxia are summarized in Table 1.

**Table 1:** Transcript variants of human and murine Naa10 implicated in hypoxia. Summary of the Naa10 isoforms that were used in hypoxia research including their accession number and corresponding citation. Two other isoforms, hNaa10^131^ and mNaa10^198^ (Kim et al., 2006), and a predicted isoform X1 hNaa10^184^ (XM_005277911), have not been included in this table.

| name | Gene bank accession no + isoform |  | Citation |
| --- | --- | --- | --- |
| mNaa10<sup>235</sup> | NM_019870 | transcript variant 1 | (Sugiura et al., 2003; Chun et al., 2007; Seo et al., 2014) |
| mNaa10<sup>225</sup> | BC027219 | transcript variant 2 | (Jeong et al., 2002; Yoo et al., 2006; Chun et al., 2007; Lee et al., 2010b; Seo et al., 2014) |
| hNaa10<sup>235</sup> | NM_003491 | transcript variant 1 | (Arnesen et al., 2005b; Bilton et al., 2005; Fisher et al., 2005; Chang et al., 2006; Murray-Rust et al., 2006; Chun et al., 2007; Lim et al., 2008; Seo et al., 2014) |
| hNaa10<sup>220</sup> | NM_001256119 | transcript variant 2 | - |
| hNaa10<sup>229</sup> | NM_001256120 | transcript variant 3 | - |

The C-terminal domain (aa 158-225) of mouse Naa10^225^ completely differs from mNaa10^235^ and hNaa10^235^ (Kim et al., 2006), whereas mNaa10^235^ and hNaa10^235^ share 96 % sequence similarity. All isoforms share an N-terminal N-Acyltransferase superfamily domain. Transfection of mNaa10^225^ into HeLa cells strongly decreased the HIF-1α protein and VEGF mRNA levels under hypoxia whereas both other variants had only minor effects (Kim et al., 2006; Seo et al., 2014). In immunoprecipitates isolated by an acetyl-lysine antibody from HeLa cells treated with the proteasome inhibitor MG132, HIF-1α could be detected when the cells were transfected with mNaa10^225^ but was nearly undetectable when the cells were transfected with mNaa10^235^ or hNaa10^235^ (Kim et al., 2006). The authors conclude that mNaa10^225^ increased the level of HIF-1α acetylation under normoxic conditions whereas the two other analyzed variants had only weak effects. However, it still cannot be excluded that in these experiments another, yet not identified, acetylated protein co-precipitates with HIF-1α. Moreover, it is not clear whether Naa10 directly acetylates HIF-1α or if another acetyl transferase is involved that is stimulated by Naa10. The acetyltransferase activity of Naa10 seems to be important for its action in hypoxia since mutated variants of mNaa10^225^ which are not able to undergo autoacetylation (K136R) or acetyl-CoA binding (K82A/Y122F) were unable to bind to HIF-1α and did not increase HIF-1α acetylation in HeLa cells as detected by acetyl-lysine antibodies (Seo et al., 2014).

The expression pattern of Naa10 isoforms was analyzed by western blotting in the human cervical adenocarcinoma HeLa, fibrosarcoma HT1080, in the lung adenocarcinoma H1299 cell line, as well as in the murine fibroblast cell line NIH3T3. hNaa10^235^ was identified as the major form in the human cell lines, whereas mNaa10^235^ and mNaa10^225^ were both detected in the murine cell line (Kim et al., 2006), indicating, that – at least in human cells – the shorter Naa10 variant plays only a minor role in hypoxia response.

Other studies suggest that the regulation of HIF-1α by Naa10 might require other factors/regulators such as deacetylases and/or may be dependent on different signaling pathways. In this context, it has been shown that the suppression of HIF-1α by Naa10 is dependent on the expression level of MTA-1 (metastasis-associated protein 1) a component of the nucleosome remodeling and histone deacetylase (HDAC; NuRD) complex. Upon CoCl_2_-induced hypoxia, MTA-1 is expressed, binds to the ODD and C-terminal domain of HIF-1α, stimulates deacetylation of HIF-1α at Lys532, thereby stabilizing HIF-1α and enhancing its interaction with HDAC1 leading to increased transcriptional activity (Yoo et al., 2006). Trichostatin (TSA), a potent specific inhibitor of HDAC, increases acetylation of HIF-1α and decreases HIF-1α stability (Yoo et al., 2006). Therefore the authors speculate that MTA-1 may counteract Naa10 in the regulation of HIF-1α stability by activating HDAC1. In contrast to this, it has been shown that TSA does not induce acetylation or regulate stability of HIF-1α but induces hyperacetylation of p300 thereby reducing the interaction of p300 with HIF-1α repressing the transactivation potential of HIF-α/p300 complex in HeLa cells (Fath et al., 2006). Similarly, it could be shown that Naa10 transfection had no effect on hypoxia-induced stabilization of HIF-1α in HeLa, HepG2 and MCF-7 cells that express high levels of MTA-1, but suppresses stabilization in HEK293 cells (expressing low levels of MTA-1) (Yoo et al., 2006). Under normoxic conditions, MTA-1 expression is repressed and overexpressed HIF-1α is proteasomally degraded upon Naa10 transfection even in MCF-7 cells (Yoo et al., 2006). These findings indicate that the expression level of MTA-1 as well as cellular oxygen status might impact Naa10 function in regulating HIF-1α signaling. The pro-fibrotic connective tissue growth factor (CTGF) can also regulate angiogenesis by interacting with VEGF. Transfection of the human lung adenocarcinoma cells CL1-5 and A549 with CTGF leads to increased Naa10 expression levels, increased HIF-1α acetylation, enhanced interaction and ubiquitination with/by VHL and accelerated the proteasomal degradation of HIF-1α (Chang et al., 2006). Additionally, knockdown of Naa10 with antisense decreased the level of acetylated HIF-1α and restored VEGF-A expression in these cells (Chang et al., 2006). Therefore the authors hypothesize that the Naa10-dependent regulation of HIF-1α might be dependent on CTGF and speculate that certain cellular proteins are induced by CTGF that work coordinately with Naa10 to affect HIF-1α protein stability.

A different study showed that Naa10^235^ regulates the response to hypoxia through a different pathway, suggesting a crosstalk between HIF-1α and the Wnt-signaling pathway. In cells with activated Wnt-signaling, HIF-1α competitively dissociates Naa10 from β-catenin preventing its acetylation under hypoxic conditions (Lim et al., 2008). This in turn represses β-catenin/TCF4 transcriptional activity, resulting in c-Myc suppression and p21(cip1) induction and proliferation inhibition (Lim et al., 2008). In this context, studies from APC^Min/+^ mice are very interesting. These mice harbor a Min (multiple intestinal neoplasia) mutation leading to a truncated version of APC (adenomatous polyposis coli), a component of the β-catenin-destruction complex. Whereas the APC knockout is embryonic lethal, heterozygous mice expressing this variant are viable but spontaneously develop intestinal polyps. Apc^Min/+^/mNaa10^225^ mice (generated by crossbreeding Apc^Min/+^ and mice expressing mNaa10^225^ from the actin promoter) are characterized by a decreased polyp size and number, and the tumors contain a decreased VEGF-A level and microvessel density compared to Apc^Min/+^ mice (Lee et al., 2010b). Furthermore, expression of mNaa10^225^ in B16F10 and MKN74 reduced endogenous HIF-1α protein and VEGF-A mRNA and protein levels under hypoxia and suppressed migration, whereas co-expression of HIF-1α K532 mutant abolished these effects (Lee et al., 2010b).

Somewhat related to this is a study that analyzed Naa10 regulation in oxidative stress response. Under normoxic conditions, reactive oxygen species (ROS) are inevitable generated and enzymes have evolved to counteract these metabolites. One such enzyme is the methionine sulfoxide reductase (MSR) that consists of MSRA (methionine sulfoxide reductase A) and MSRB (methionine sulfoxide reductase B) that reduce S-sulfoxide and R-sulfoxide, respectively. Naa10 interacts with MSRA and acetylates it on K49 thereby inhibiting MSRA activity as shown in *in vitro* assays (Shin et al., 2014). Furthermore, Naa10 overexpression in A549 and H1299 cells or transgenic mouse (overexpressing Naa10 in kidney and liver) reduced MSRA activity as measured by an increase of ROS, thereby promoting cell death (Shin et al., 2014). The authors speculate that Naa10 regulation in hypoxia might be related to its role in oxidative stress, as intracellular ROS increases during hypoxia; however, the mechanism remains unclear.

**Figure 4:**
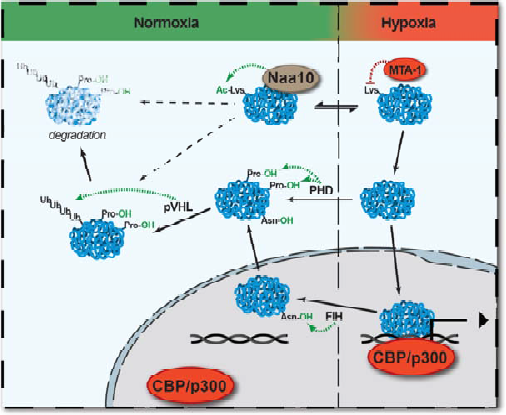
Possible role of Naa10 in cellular hypoxia. Under normoxic conditions, HIF1α is hydroxylated by PHD proteins on prolines, leading to the subsequent ubiquitination by pVHL and rapid proteasomal degradation of HIF1α. Furthermore, hydroxylation of Asn_903_ by FIH disrupts binding of HIF1α to its transcriptional co-activators p300/CBP, and hence the transcriptional activity of the HIF-1α. Many reports have shown a direct interaction of Naa10 with HIF-1α, suggesting a link in this pathway. Naa10 might either directly or indirectly stimulate HIF1α lysine acetylation, thereby destabilizing HIF1α, possibly through promoting its interaction with pVHL and PHDs. MTA1 counteracts the action of Naa10 by recruiting HDAC1 which leads to a deacetylation and subsequent stabilization of HIF-1α.

Although the direct interaction of Naa10 and HIF-1α is widely accepted and indicates a link between Naa10 and hypoxia, the physiological significance of Naa10 in this context is still unclear (**Error! Reference source not found.**). It seems quite established that HIF-1α acetylation at K532 is a destabilizing factor in normoxia. However, further studies have to address the following questions:

- Is HIF-1α K532 directly acetylated by Naa10? Many studies have questioned this idea and the use of anti-lysine-antibodies is prone to misinterpretation as the position of acetylation is not defined. p300 itself for example acetylates HIF-1α at K709 thereby regulating its stability (Geng et al., 2012) and changes in the acetylation status on this position would affect the above assay.
- If Naa10 does not directly acetylate HIF-1α K532, does it promote its acetylation indirectly? If so, what is the unknown enzyme and how does Naa10 regulate its target recognition/catalytic activity?
- What effect does the cellular status have on Naa10 function in normoxic/hypoxic conditions? Several studies show that Naa10 function in hypoxia depends on other factors including MTA-1, CTGF and β-catenin. The activation of some of these pathways in different model systems could explain some of the observed discrepancies. For example, MTA-1 expression strongly correlates with hypoxia and some cells only weakly express MTA-1 under normoxic conditions (Yoo et al., 2006). Additionally, HIF-1α and β-catenin are deregulated in many cancers (Zhong et al., 1999; Talks et al., 2000; Clevers and Nusse, 2012) and therefore might be deregulated in many cancer-derived cell lines. How does that influence the action of Naa10 in hypoxia? Therefore, the choice of the cell system to study might strongly influence the outcome of Naa10 knockdown or overexpression.
- What are the isoform-specific Naa10 effects and what are the differences between species? In mouse, mNaa10^225^ had a strong effect on HIF-1α acetylation and stability, whereas mNaa10^235^ and hNaa10^235^ had only minor effects. However, all isoforms interact with HIF-1α and possess an N-terminal aceyltransferase domain. Therefore the differences in the regulation remain unclear. Furthermore, unlike in mouse, in human cell lines, mainly Naa10^235^ is expressed, which seems to differ in its functionality from mNaa10^225^ (Kim et al., 2006).

#### 2.1.4 Bone formation

Phenotypic analyses from transgenic mice implicate Naa10 in bone development and link Naa10 to the runt-related transcription factor 2 (Runx2). In the early phase of bone development, Runx2 regulates osteoblast differentiation through transactivation of many genes including collagen α1, osteopontin, bone sialoprotein (BSP) and osteocalcin (OCN). A recent study showed that Naa10 interacts with the RUNT domain of Runx2, acetylates it at Lys225, and disrupts Runx2/CBFβ interaction, thereby inhibiting Runx2 transcriptional activity as shown in *in vitro* association, acetylation and reporter gene assays, respectively (Yoon et al., 2014). Additionally, knockdown of Naa10 augmented the stimulatory effects of BMP-2 on osteoblastogenesis in a rat calvarial critical-size defect model and enhanced the differentiation in primary mouse osteoblasts (Yoon et al., 2014). Furthermore, overexpression of Naa10 in transgenic mice resulted in calvarial fontanels being less closed and bones being less dense, a lower osteoblast surface and reduced mRNA of osteoblastic genes (delayed bone development), whereas Naa10 knockout mice exhibit normal skeletal structure at day 3 after birth and bones developed to a greater extent, supporting the idea that Naa10 functions in Runx2-mediated osteogenesis *in vivo* (Yoon et al., 2014).

Naa15 also has been implicated in osteogenesis. Studies of the osteocalcin promoter identified Naa15, Ku70 and Ku80 as possible regulators. These proteins bind the osteocalcin promoter and activate osteocalcin expression synergistically with Runx2, suggesting a functional interactions between Ku, Naa15, and Runx2 (Willis et al., 2002). In contrast to the above, Yoon and colleagues showed that knockdown of Naa15 does not influence BMP-2 induced osteoblast differentiation and does not affect the Naa10-dependent acetylation of Runx2 (Yoon et al., 2014).

#### 2.1.5 Neuronal development

An early study demonstrated that mNaa15 expression is temporally regulated in mouse brains during development (Sugiura et al., 2001). In a follow up study, *in situ* hybridization showed that mNaa10 and mNaa15 are highly expressed throughout mouse brain development in areas of cell division and migration, but are down-regulated as neurons differentiate and mitotic and migratory activities subside (Sugiura et al., 2003). During mouse brain development, Naa10 and Naa15 levels stay high in the olfactory bulb, neocortex, ventricular zone and hippocampus, regions where cell division and migration are still prominent in the neonate, suggesting that NatA plays an important role in the generation and differentiation of neurons (Sugiura et al., 2003). In line with that, it could be shown that the expression of Naa10 and Naa15 increases during neuronal dendritic development of cerebellar Purkinje cells, and knockdown of Naa10 significantly limited the dendritic extension of cultured neurons (Ohkawa et al., 2008). Furthermore, both proteins colocalize with microtubules (MTs) in dendrites, Naa10 rescues an arborization defect of HDAC6-(a major deacetylases of α-tubulin)-overexpressing cells, and an immunopurified NatA complex acetylates purified α-tubulin (Ohkawa et al., 2008). Tubulin acetylation is involved in neuron polarization, neurite branching and promotes vesicular transport on MTs in differentiated neurons (Perdiz et al., 2011). Therefore the authors speculate that Naa10 counteracts HDAC6 by acetylating α-tubulin, thereby promoting MT stability for dendritic development. A different study suggests that HDAC6 reverses Naa10-mediated acetylation of Runx2, suggesting that HDAC6 might be a major counter player of Naa10 (Yoon et al., 2014)

However, other studies have shown that αTat1 (α-tubulin K40 acetyltransferase) is a predominant/major α-tubulin acetyltransferases *in vivo* in mammals and nematodes and finetunes hippocampus development (Shida et al., 2010; Kim et al., 2013). As a side note, in *D. melanogaster*, Naa50 has been shown to acetylate β-tubulin at lysine252 on soluble tubulin heterodimers but not tubulins in MTs. thereby modulating MT polymerization (Chu et al., 2011). More studies are needed to elucidate the mechanism for how Naa10 regulates neuronal differentiation and what the involved substrates are.

#### 2.1.6 Naa10 in human disease

As we will discuss in greater detail below, the links between Naa10 and diseases in humans are only recently emerging. For example, many biochemical and cellular studies link Naa10 to neurodegenerative disorders, such as Alzheimer’s, Parkinson’s and Huntington’s disease, where abnormal manifestations of single proteins are believed to contribute to disease. Recently, several studies identified mutations of *NAA10* in human patients with discrete symptoms, including an aged appearance, craniofacial anomalies, hypotonia, developmental delay, growth retardation and Lenz microphthalmia syndrome. However, the small phenotypic overlap between these patients suggests that different substrates or functions of Naa10 might be affected.

In the case of Alzheimer’s disease, it has been shown that Naa10 binds via its 50 C-terminal amino acids to the cytoplasmic domain of the type I membrane protein amyloid precursor protein (APP) APP (β-amyloid precursor protein) and colocalizes with it in HEK293 cells (Asaumi et al., 2005). In Alzheimer’s disease, APP gets processed into beta-amyloid (Aβ) fragments that accumulate into plaques of abnormally folded proteins in the brain. Overexpression of the NatA complex (Naa10 wild type/Naa15 wild type) in HEK293 cells decreases endocytosis of APP and suppresses Aβ40 secretion, whereas the expression of an enzymatic-dead NatA (Naa10 R82A or G85A/Naa15 wild type) attenuated this suppression (Asaumi et al., 2005). However, the mechanism by which NatA regulates Aβ production remains unclear. Neither a direct acetylation of APP nor any other of the involved proteins have been shown, and it is noteworthy that the N-terminus of APP faces outside of the cell.

Aside from that, it has been shown that the natural C-terminal fragments of Tau (another Alzheimer’s disease-associated protein), and α-synuclein (associated with Parkinson’s disease) are NTA, and the sequence properties point to NatB as the corresponding NAT. Indeed, NatB has been identified in a genetic screen for deletion mutants with mislocalisation of α-synuclein, and the knockout of NatB diminishes the membrane localization of α-synuclein in yeast (Zabrocki et al., 2008). Furthermore, NTA of α-synuclein has been shown to promote its soluble, intrinsically alpha-helical structure, thereby inhibiting aggregation (Kang et al., 2012; Kang et al., 2013), promoting membrane/vesicle binding (Maltsev et al., 2012; Bartels et al., 2014) and interaction with calmodulin (Gruschus et al., 2013). Another group showed that NTA has no major influence on α-synuclein aggregation, synaptosomal membrane binding and only slightly increased helicity of the N-terminal region as shown by NMR, static light scattering, and sedimentation assays on recombinantly expressed α-synuclein (Fauvet et al., 2012). Proteofection of these α-synuclein variants into HeLa cells revealed that NTA does not affect subcellular localization or membrane binding capacity of α-synuclein (Fauvet et al., 2012) and no differences in the acetylation state for α-synuclein in control or PD/DLB brain samples were found (Ohrfelt et al., 2011). These studies indicate that NTA of α-synuclein either promotes the soluble state or has a very weak effect or no effect at all on α-synuclein structure and solubility,; however, another group has shown that NTA rather promotes α-synuclein oligomer formation (Trexler and Rhoades, 2012). Future work has to resolve these discrepancies and elucidate whether NTA regulates physiological dimer formation or pathological aggregation of α-synuclein. For a recent review on this topic see (Alderson and Markley, 2013).

The identification of HYPK as a stable interaction partner of NatA provides the third link to neurodegenerative disorders. HYPK acts as a chaperone and reduces Htt (Huntingtin) polyglutamine (polyQ) aggregation, which is believed to cause Huntingtin’s Disease (Raychaudhuri et al., 2008). Knockdown of both Naa10 and Naa15 decreased HYPK protein levels and knockdown of HYPK or Naa10 resulted in increased aggregation of an Htt-reporter construct, suggesting that both HYPK and NatA prevent Htt aggregation (Arnesen et al., 2010). The mechanism of this has not been identified.

In addition, NatA function has been shown to regulate protein folding through the actions of chaperones, thereby promoting prion [*PSI*^+^] propagation in yeast (see below), illustrating the important contributions of NTA to amyloidogenesis and its biological consequences (Holmes et al., 2014).

Recently, two families with a lethal X-linked disorder of infancy were described, with this new syndrome being named Ogden syndrome in honor of where the first family lives (in Ogden, Utah) (Figure 5). The disorder comprises a distinct combination of an aged appearance, craniofacial anomalies, hypotonia, global developmental delays, cryptorchidism, and cardiac arrhythmias (Rope et al., 2011). Using X chromosome exon sequencing and a recently developed variant annotation, analysis, and selection tool (VAAST), a c.109T>C (p.Ser37Pro) variant in *NAA10* was identified (Rope et al., 2011). Functional analysis of this mutation revealed a decreased enzymatic activity of the S37P mutant in *in vitro* acetylation assays towards canonical (NatA) and new (likely monomeric Naa10) substrates (Rope et al., 2011), and further characterization of the catalytic activity (k_cat_/K_m_) indicated that the S37P mutation possibly alters peptide substrate binding or release without affecting acetyl-CoA binding (Myklebust et al., 2014). In agreement with a diminished catalytic activity of the Naa10 S37P mutant, quantitative N-acetylome analyses (COFRADIC/MS) showed a reduced degree of NTA for a majority of NatA substrates in a yeast Ogden syndrome model (Van Damme et al., 2014). These cells overexpress either the human wt (hNaa15/hNaa10 wild type) or mutant NatA (hNaa15/hNaa10-S37P) complex in the yNatA knockout background (y*NAA10*Δ/y*NAA15*Δ). Similarly, quantitative analysis of proteome wide acetylation in B-cells and fibroblasts from the Ogden families revealed a reduction of the *in vivo* acetylation status for a small subset of N-termini of classical NatA substrates and some NatE substrates (Myklebust et al., 2014). Co-immunoprecipitation experiments showed that the S37P Ogden mutation also decreased the interaction of Naa10 with Naa50, the catalytic subunit of NatE, providing a possible explanation for the latter finding (Myklebust et al., 2014). In yeast, immunoprecipitation assays from the Ogden model strains with an anti-Naa15 antibody showed a 2 fold reduction in NatA complex formation and a reduced acetyltransferase activity of the mutated Naa10 compared to wild type (Van Damme et al., 2014). In agreement with that, immunoprecipitation experiments in HEK293 cells and patient fibroblasts showed a reduced NatA complex formation, disruption of Naa10-Naa50 interaction, and impaired enzymatic activity of Naa10-S37P (Myklebust et al., 2014). Structural dynamic simulations of both, wild-type human NatA complex and the S37P mutant, showed perturbations of the hNaa10-hNaa15 interaction interface and changes in the flexibility of key regions for substrate binding which would explain the impaired Nt-acetyltransferase activity and complex stability of NatA-S37P (Myklebust et al., 2014). Taken together, these data suggest a diminished functionality of hNatA S37P *in vitro* and *in vivo.*

**Figure 5:** Pictures related to Ogden Syndrome. Shown are pictures of individual III:4 and III:6 from family one (A) and individual II:1 and individual III:2 from family 2 (B).

*** Photos will appear in the published book chapter. Medicolegal reasons prevent their publication as part of this BioRxiv preprint.

The functional consequences of the S37P mutation were studied in various models. In the recently developed Ogden syndrome heterologous yeast model, human wild type NatA rescued the sensitivity of the yNatA knockout strain towards caffeine and cycloheximide, whereas the S37P mutant was qualitatively shown to only partially rescue the phenotypes in yeast strain spotting experiments (Van Damme et al., 2014), indicating an increased sensitivity of the S37P cells towards stress. Fibroblasts from Ogden patients displayed altered morphology, growth and migration characteristics, possibly linked to a perturbed retinoblastoma (Rb) pathway, as Rb1 proteins levels were found to be increased; however, Naa10 protein levels or subcellular localization of Naa10 were unaffected in B-cells and fibroblast from Ogden males (Myklebust et al., 2014). These cellular studies indicate that the Ogden mutation induces changes in general cellular key features, including proliferation, and sensitizes cells towards stress. Analysis of the X-chromosome inactivation pattern of the Ogden family revealed skewing of the affected allele in fibroblasts from carrier women, indicating a decreased fitness or production rate of these cells (Myklebust et al., 2014).

The mechanism for how reduced NTA acetylation of a subset of NatA substrates could induce the severe phenotype observed in Ogden syndrome is still unknown. However, now that some putative affected proteins are identified, further studies have to elucidate the specific consequences of a decrease in acetylation on the function of these proteins. In a first attempt, analysis of the stability of 8 affected proteins (reduced acetylation status) revealed no change of the steady state protein levels except for THOC7 (THO complex subunit 7 homolog), and mutations of THOC7 (G2V or G2P) that partially or completely abrogated NTA, respectively, strongly reduced steady-state protein level in A431cells, which could be rescued upon proteasomal inhibition (Myklebust et al., 2014). Similarly, siRNA-mediated knockdown of NatA decreased the half-life of THOC7 in cycloheximide chase experiments, indicating that NTA confers stability on THOC7 (Myklebust et al., 2014), which possibly contributes to the Ogden phenotype. However, further studies are necessary to affirm this assumption and analyze the effects of NTA-impairment on affected proteins.

Another group used exome sequencing to identify a R116W mutation in a boy and a V107F mutation in a girl in two unrelated families with sporadic cases of non-syndromic intellectual disabilities, postnatal growth failure and skeletal anomalies (Rauch et al., 2012; Popp et al., 2014). A relatively mild phenotype in the boy compared to a strong defect in the girl correlates with the remaining catalytic activity of Naa10 as measured in *in vitro* NTA assays, suggesting that the “N-terminal acetyltransferase deficiency is clinically heterogeneous with the overall catalytic activity determining the phenotypic severity” (Popp et al., 2014). In a study from 2013 where 106 genes proposed to be involved in monogenic forms of X-linked intellectual disability (XLID) were reassessed using data from the National Heart, Lung, and Blood (NHLBI) Exome Sequencing Project, this mutation in Naa10 reported in 2012 was marked as “awaiting replication” with at least one other family (Piton et al., 2013).

Another recent study implicated a mutation in *NAA10* in a single family with Lenz microphthalmia syndrome (LMS), a very rare, genetically heterogeneous X-linked recessive disorder characterized by microphthalmia or anophthalmia, developmental delay, intellectual disability, skeletal abnormalities and malformations of teeth, fingers and toes. Exome sequencing in this family with three affected brothers of LMS identified a splice mutation in the intron 7 splice donor site (c.471+2T→A) of *NAA10* (Esmailpour et al., 2014). Patient fibroblasts lacked expression of full length Naa10 protein and displayed cell proliferation defects, although it was not clear from the study if the control (wild type) cell lines were taken from unaffected family members or from completely unrelated individuals. STRA6, a retinol binding protein receptor that mediates cellular uptake of retinol/vitamin A and plays a major role in regulating the retinoic acid signaling pathway, was found to be dysregulated, as shown by expression array studies, and retinol uptake was decreased in patient cells (Esmailpour et al., 2014). Finding additional unrelated families with the exact same mutation, or at least other mutations in *NAA10* and also with this similar phenotype, will be very important, particularly as there is very little overlap between Ogden Syndrome and this phenotype. The field of human genetics has begun to coalesce around the notion that mutations should be found in more than one family prior to making strong assertions that any reported mutation really contributes to any particular phenotype being studied. In this regard, the two unrelated families with Ogden Syndrome were both found to have the exact same mutation, with perfect segregation in 5 affected boys in one family and in 3 affected boys in the other family, alongside not being found in several unaffected males in at least one of the families. However, even though this was reported in 2011, there have not yet been any reports of even a third family with the exact same mutation, thus highlighting the major issues with finding additional families, just as what appears to have occurred with these above singleton families.

One problem is that the phenotypic changes that can be observed with NatA mutations might arise from the reduced acetylation status of multiple target proteins. A similar phenomenon has been described for the mating type switching (Geissenhöner et al., 2004; Wang et al., 2004) and the [*PSI+*] prion phenotype (Holmes et al., 2014) in *S. cerevisiae* (see below), plus there are many tissue-specific effects of the Ogden Syndrome and other mutations. Therefore, the observed differences in phenotype might result from differences in the acetylation status of multiple proteins and/or protein complexes in different tissues and/or in combination with global changes in cellular performance, e.g. induced by acetylation-dependent changes in protein biogenesis. This shows the inevitable need to study the effects of NTA on individual proteins as well as on a more general level.

### 2.2 Function of Naa15/Naa16/Tubedown

In contrast to Naa10, the auxiliary subunit of the NatA complex, Naa15, is less intensively studied. However, recent evidence suggests that Naa15 is involved in cell survival and tumor progression. Quantitative analysis of Naa15 expression in neuroblastic tumors (neuroblastomas, ganglioneuroblastomas, and ganglioneuromas) and normal adrenal tissues revealed that Naa15 is highly expressed in tumors with significant neuroblastic components, unfavorable histopathology, advanced stage, high-risk group, and poor outcome. Additionally, retinoic acid-induced neuronal differentiation responses of neuroblastoma cells were associated with a significant decrease in Naa15 expression, whereas limited neuronal differentiation responses to retinoic acid were associated with little or no decrease in Naa15 expression, indicating that Naa15 expression is linked to the differentiation status and aggressiveness of neuroblastic tumors (Martin et al., 2007). Specific knockdown of hNAA16 in HEK293 cells induces cell death, suggesting an essential role for hNaa16p in human cells (Arnesen et al., 2009a). Furthermore, Naa15 is overexpressed in tumor cells in PTC (papillary thyroid carcinomas) and especially in a Burkitt lymphoma cell line indicating a function in the progression of papillary thyroid carcinomas, but heterologous expression of Naa15 did not alter the cellular proliferation rate (Fluge et al., 2002).

Recent studies suggest that Naa15 may play a role in the maintenance of a healthy retina. The expression of Naa15 during oxygen-induced retinal neovascularization in mice and in a specimen of stage 3 human retinopathy of prematurity (IHC) is suppressed, indicating that loss of retinal endothelial Naa15 expression is associated with retinal fibrovascular growth and thickening (Gendron et al., 2006). In line with this, Naa15 loss resulting from old age or conditional knockdown was associated with retinal lesions showing significant extravascularly localized albumin and correlated with increased activity of senescence (Gendron et al., 2010). Furthermore, in endothelial-specific-Naa15-knockdown mice a significant leakage of albumin from retinal blood vessels compared with control age-matched mice was observed contributing to the retinal pathology (Paradis et al., 2008). Naa15 interacts with the actin binding protein cortactin in cytoplasmic regions at the cortex of cultured endothelial cells thereby regulation endothelial cell permeability (Paradis et al., 2008).

### 2.3 Function of Naa50

In yeast (*S. cerevesiae*), deletion of the N^α^-terminal acetyltransferase Naa50 does not cause a detectable phenotype (Gautschi et al., 2003). However, early studies in Drosophila showed that Naa50 is required for establishing sister chromatid cohesion. Mutation of *Drosophila melanogaster* Naa50 (san/separation anxiety) disrupts centromeric sister chromatid cohesion, activates the spindle checkpoint, and causes metaphase delay or arrest (Williams et al., 2003). Recent observations confirmed that disruption of Naa50 leads to severe chromosome segregation and chromosome resolution/condensation defects during mitosis. These defects seem to be specific for somatic cells, since mitotic function was not affected in female germ line stem cells during oogenesis (Pimenta-Marques et al., 2008).

This function seems to be conserved in metazoans as indicated in a later study, where enzymatic active Naa50 was critical for sister chromatid cohesion in HeLa cells, independently of NatA, as shown in rescue experiments with a catalytic-defective Y124F mutant (Hou et al., 2007). However, the mechanism by which Naa50 regulates sister chromatid cohesion has not been elucidated. One possible substrate that matches the preferred N-terminal sequence for the NAT activity of hNaa50 is TIMELESS-interacting protein, a protein that is reportedly involved in chromatid cohesion (Evjenth et al., 2009). But, we have not found any published experimental data regarding this yet.

Recently, Naa50 was described to have β-tubulin acetyltransferase activity. Dynamic instability is a critical property of microtubules (MTs). Naa50 was reported to acetylate β-tubulin at a novel site (lysine 252) on soluble tubulin heterodimers but not on polymerized tubulins in MTs. After cold-induced catastrophe, MT regrowth was accelerated in Naa50-siRNA knockdown cells while the incorporation of an acetylation-mimicking tubulin K252A/Q mutant into MTs was severely impeded (Chu et al., 2011). Because K525 interacts with the α-tubulin-bound GTP, the authors propose that acetylation slows down tubulin incorporation into MTs by neutralizing the positive charge on K252, which allows tubulin heterodimers to form a conformation that disfavors tubulin incorporation.

## 3 NatA in other organisms

Since its first identification in a yeast screen (Whiteway and Szostak, 1985) homologues for Naa10 have been found in a variety of other organisms. In an early review, Polevoda and Sherman used the general BLAST server from the National Center for Biotechnology Information (NCBI) to identify orthologs of yeast NATs. They found candidates for Naa10 in *S. pomb*e, *C. elegans*, *D. melanogaster*, *A. thaliana*, *T. brucei*, *D. discoideum*, *M. musculus* and *H. sapiens,* and orthologs for Naa50 were found in *S. pombe*, *C. elegans*, D*. melanogaster*, *A. thaliana* and *H. sapiens* (Polevoda and Sherman, 2003b). Since then, many orthologs of Naa10 have been identified in a variety of different organisms, showing the generality of the NTA machinery in eukaryotes. Also, a proteomic comparison of acetylated proteins in *S. cerevisiae* and human HeLa cells revealed that the NatA substrate specificity is almost the same in both organisms indicating that the acetylation of proteins is evolutionarily conserved (Arnesen et al., 2009b). In the archeae *S. solfataricus*, only a single ssNaa10 has been identified to date with a more relaxed substrate specificity (acetylating NatA, NatB and NatC-type substrates) (Mackay et al., 2007). A recent structural comparison of ssNaa10 with *S. pombe* Naa10 showed that the active site of ssNaa10 represents a hybrid of Naa10 and Naa50 and, as a result, can facilitate acetylation of Met- and non-Met-containing amino-terminal substrate peptides (Liszczak and Marmorstein, 2013). Therefore, the authors suggest that this ssNaa10 is an ancestral NAT variant from which the eukaryotic NAT machinery evolved (Liszczak and Marmorstein, 2013).

Naa10 homologues have also been identified in the protozoan parasites *Trypanosoma brucei* and *Plasmodium falciparum* (Ingram et al., 2000; Chang et al., 2008). In *T. brucei*, disruption of both alleles was lethal, indicating that Naa10 is essential.

### 3.1 Naa10 in *C. elegans*

The *C. elegans* ortholog to Naa10 (abnormal dauer formation-31, DAF-31, K07H8.3) was first identified in a genetic screen to identify novel mutants affecting dauer and dauer-like larvae (Chen et al., 2014). In favorable environments, the embryonic development in *C. elegans* consists of four larval stages (L1-L4); however, when food is limited, the nematodes arrest at the L1 dauer diapause, a long-lived stress resistant stage (Fielenbach and Antebi, 2008). The insulin/IGF-1 signaling pathway regulates, among others, dauer and stress resistance in *C. elegans*. Activation of the insulin/IGF-1 transmembrane receptor (IGFR) ortholog DAF-2 results in the activation of a signaling cascade that eventually inhibits the DAF-16/FOXO (forkhead box O transcription factor) transcription factor, by promoting its sequestration in the cytoplasm, thereby negatively regulating dauer formation, longevity, fat metabolism and stress response (Murphy and Hu, 2013). The identified mutant animals (bearing a 393 bp deletion in the promoter and N-terminal region of *NAA10/daf-31*) formed dauer-like uncoordinated (Unc) larvae in the presence of dauer-inducing pheromones and under starvation, but unlike wild type animals, were sensitive to SDS and did not resume development when food was provided (Chen et al., 2014). Furthermore, overexpression of Naa10 stimulated transcriptional activity of DAF-16 thereby enhancing the longevity phenotype of DAF-2/insulin receptor mutants, indicating that Naa10 regulates *C. elegans* larval development dependent on the DAF-2/IGFR pathway (Chen et al., 2014). However, the exact mechanism remains unclear and the sequence of DAF-16 (starting with MQLE…) indicates that it is not a direct target for NTA by Naa10. In parallel with the above, another report was published implicating NatC in the regulation of stress resistance towards a broad-spectrum of physiologic stressors, including multiple metals, heat, and oxidation. Particularly, Naa35 (NATC-1, NatC auxiliary subunit, T23B12.4) is transcriptionally repressed by DAF-16 and mutations in *NAA35* or *NAA30* increased dauer formation, indicating that NatC is an effector of the insulin/IGF-1 signaling pathway downstream of DAF-16 (Warnhoff et al., 2014). Taken together, these data shows that NTA may play an important role in *C. elegans* development at different levels, although the exact mechanism for this action remains elusive. The ortholog of *NAA15* in *C. elegans*, *hpo-29*, was identified in an RNAi screen for genes that, when knocked down, result in hypersensitivity towards pore-forming toxins (Kao et al., 2011). However, the mechanism of this action was not analyzed.

### 3.2 NatA in Yeast (*S. cerevisiae* unless indicated otherwise)

The yeast *NAA10* gene was first identified in a genetic screen searching for mutants with an arrest-defective phenotype (mutants that exhibit a high proportion of budding cells and are resistant to mating α-factor) and therefore named *ARD1* (arrest-defective) (Whiteway and Szostak, 1985). Later studies showed that *NAA15* knockout cells exhibit the same phenotype as a *NAA10* knockout: temperature sensitivity, a growth defect, de-repression of the silent mating type locus (*HML*), and failure to enter G_o_ phase (Lee et al., 1989; Mullen et al., 1989), highlighting that both Naa10 and Naa15 function together to regulate gene silencing and mating in yeast. A later study found that disruption of NatA elevated protein aggregation and modulated the expression as well as the recruitment of molecular chaperones, including Hsp104 and Hsp70, to protein aggregates, potentially contributing to the observed heat sensitivity (Holmes et al., 2014). In agreement with that, overexpression of the Hsp70 protein Ssb1 partially suppressed the temperature sensitivity and derepression of the silent mating type loci caused by *NAA15*Δ or *NAA10*Δ (Gautschi et al., 2003). Furthermore, disruption of NatA (Naa10 or Naa15) affects general cell fitness as assayed by the ability of cells to restart their cell cycle when nutrients are added after starvation (Aragon et al., 2008; van Deventer et al., 2014).

#### 3.2.1 Mating type/silencing defect

In yeast, sexual reproduction is regulated by nonhomologous alleles, the mating type. The alleles of the mating type locus, *MAT**a*** and *MAT*α, encode for proteins specifically regulating the expression of sex specific genes. Two silent loci, *HML* and *HMR,* are located on opposite ends of the same chromosome-encoding *MAT* and serve as donors during the recombinational process of mating type switching. In most strains, each *HML* and *HMR* contain a cryptic copy of the mating type sequences *MAT*α and *MAT**a***, respectively (Haber, 2012). Silencing of the *HML* and *HMR* loci, rDNA loci and telomeres is accomplished by trans-acting factors like Sir (silent information regulator) proteins that bind to specific silencer sequences, called *HML*-E, *HML*-I, *HMR*-E and *HMR*-I (Haber, 2012). Orc1p, the large subunit of the origin recognition complex (ORC), binds to the silencer and recruits Sir1p (Fox et al., 1997). The N-terminal domain of Orc1p is related to Sir3p and is required for mating type silencing (Bell et al., 1995). Binding of these proteins at the silencer region recruits the Sir2p/Sir3p/Sir4p complex. The histone deacetylase Sir2p deacetylates the tails of histone H3 and H4, establishing binding sites for Sir3p/Sir4p (they bind deacetylated tails of histones) (Steglich et al., 2013). This is thought to start a sequential recruitment of additional Sir proteins, leading to the deacetylation of further histones and spreading of Sir proteins along the silencer to repress transcription in this region (Rusche et al., 2003). Similarly, silencing adjacent to telomeres is regulated in yeast, involving many of the components of the *HML/HMR* silencing machinery; however, telomeric silencing is less robust (Haber, 2012).

As mentioned before, it is well established that mutations in y*NAA10* prevent entry into stationary phase G_0_ and induce de-repression of the normally silent α information at the HML locus, leading to a mating defect in *MAT***a** cells (Whiteway and Szostak, 1985; Whiteway et al., 1987; Aparicio et al., 1991). *MAT*α *ard1* cells respond normally to **a**-factor (Whiteway and Szostak, 1985). Further studies showed that overexpression of Sir1p suppresses the mating defect of *yNAA10*Δ/*yNAA15*Δ cells (Stone et al., 1991). Comparing wild type and *NAA10*Δ mutant strains and biochemical analyses of Sir proteins revealed that NatA acetylates the N^α^-terminal alanine residues of Sir3 and of the Orc1 bromo-adjacent homology (BAH) domain but not Sir2, and this is important for the silencing of *HML* (Wang et al., 2004). Introducing either wild type Sir3 or Sir3-A2S, -A2G or -A2T mutants (all theoretically susceptible to acetylation by the NatA complex) restored the mating defect in *sir3*Δ *MAT***a** cells (Wang et al., 2004) whereas a *sir3*Δ *sir3-A2Q* mutant that abrogates acetylation by NatA had a severe defect for silencing at *HML* and telomeres (Wang et al., 2004; Ruault et al., 2011). The introduction of Sir3-A2S mutant could restore the mating of s*ir1*Δ*/sir3*Δ double knockout cells and only exhibited a slight telomeric silencing effect, whereas the Sir3-A2G or -A2T mutants all failed to mate in the absence of Sir1 (Stone et al., 2000; Wang et al., 2004; van Welsem et al., 2008). Taken together, these data suggest that mutations of the N-terminal alanine of Sir3 causes different phenotypes, ranging from a mild effect of the A2S mutation, an intermediate effect of the A2T or A2G mutations, to the A2Q mutation displaying the strongest effects. Similar effects were shown for the *MAT*α cells, although the characteristics of the mutations appears to be not as severe as for the *MAT***a** cells (Wang et al., 2004). In this context, a later study found an explanation for why NatA mutations preferentially de-repress the *HML* loci. When Sir3 becomes limiting, silencing at *HMR* is more stable than *HML*, indicating that the *HMR* locus is better able to sustain silencing at very low levels of Sir3 (Motwani et al., 2012). In agreement with that, weakening of *HMR-E* by using a synthetic variant that lacks much of the functional redundancy of the natural silencer (*HMR SS*Δ*I*) leads to a strong mating defect upon disruption of *NAA15* in *MAT*α cells (Geissenhöner et al., 2004).

An explanation for the impact of Sir3 acetylation on its function partly comes from structural studies of acetylated and non-acetylated forms of Sir3. *In vitro* studies of purified yeast chromatin fragments and Sir3 purified from wild type or *NAA10*Δ yeast showed a reduced interaction of nucleosomes with the unacetylated BAH domain (compared to the acetylated form) (Onishi et al., 2007; Sampath et al., 2009) and acetylated Sir3 formed Sir-nucleosome filaments, whereas the non-acetylated Sir3 formed short filaments with a different morphology (Onishi et al., 2007). The crystal structure of non-acetylated Sir3 indicated that the N-terminal BAH domain is disordered (Armache et al., 2011). In contrast to this, acetylated Sir3 exhibits a more structured N-terminus as demonstrated by the crystal structure of the N-terminal BAH domain of Sir3 bound to the nucleosome core particle (NCP) (Arnaudo et al., 2013). Particularly, the interaction of the N-terminal acetyl group and loop 3 of Sir3 appears to generate a rotation of the whole BAH domain toward the surface of the NCP, positioning helix 8 closer to the core histones and thereby stabilizing the interaction of Sir3 and the nucleosome (Arnaudo et al., 2013). Mutations within the N-terminal region that either prevent N-terminal methionine cleavage or replace the Ala at position 2 would disrupt this structure and reduce NPC binding as the residual initiator methionine would clash with Trp142 or long side chains would clash with the short N-terminal helix (Arnaudo et al., 2013).

Methylation of histones is another important mechanism in regulating Sir3 function/silencing. For example, methylation of H3 K79 by Dot1 abrogates binding of Sir3 to histone H3 (Onishi et al., 2007; Wang et al., 2013). Furthermore, studies showed that N^α^-terminal acetylation of Sir3 enhances its specificity for nucleosomes that are unmethylated at H3K79 (van Welsem et al., 2008). The crystal structures of the unacetylated Sir3 suggests that methylation of H3K79 decreases the potential of this lysine to form hydrogen bonds with E84 and E140 in Sir3 that would potentially result in a decreased affinity of Sir3 for the nucleosome (Armache et al., 2011). However, in the structure of the acetylated form, a set of additional interactions with the nucleosome arising from the acetylated N-terminus of Sir3 can be observed and methylation of H3K79 would create repulsion with the residues in the acetylated BAH loop 3 (Arnaudo et al., 2013), which could explain the increased affinity of acetylated Sir3 towards nucleosomes and the increased sensitivity of acetylated Sir3 towards nonmethylated nucleosomes. In contrast to this, the silencing defects of weak Sir3 mutants such as *sir3-A2G* or *sir3-A2T* mutants could be suppressed by mutations in histones H3 and H4, specifically, by H3 D77N and H4 H75Y mutations but not by knockout of Dot1, indicating that methylation is not responsible for the silencing defect of these mutants (Sampath et al., 2009).

As indicated above, another protein that has been studied, in regards to a silencing defect in NatA mutants, is Orc1. The N-terminal BAH domain of Orc1 shares 50% identity with Sir3 and the first eight residues of both proteins are completely identical (Geissenhöner et al., 2004). Furthermore, similar to Sir3, Orc1 purified from wild type yeast is found to be acetylated, but unacetylated in *NAA10*Δ or *NAA15*Δ cells (Geissenhöner et al., 2004; Wang et al., 2004). Tethering of Sir1 or Orc1 to the *HMR* silencer rescued the mating defect of *MAT*α yeast *NAA15*Δ cells, indicating that the silencing defect works upstream of Sir1 and hence, through Orc1 (Geissenhöner et al., 2004). Mutations in the penultimate residue of Orc1 (Orc1-A2P and Orc1-A2V) that inhibited its ability to be acetylated by NatA caused a severe loss of telomeric silencing; however, silencing at *HML* or *HMR* was not disrupted in these mutants (Geissenhöner et al., 2004). This is in line with the idea that *HML*/*HMR* silencing is more robust than telomeric silencing (Haber, 2012) and also suggests that the silencing phenotype of *NAA10*Δ and/or *NAA15*Δ cells is caused by an acetylation deficiency of at least two proteins, Orc1 and Sir3.

#### 3.2.2 NatA in ribosome function and protein targeting

As pointed out earlier, the NatA complex is – at least partially – associated with the ribosome. Besides that, it could also be proven that the ribosome itself is susceptible to acetylation by NATs. Edman degradation of ribosomal proteins from wt and *NAA15*Δ *S. cerevisiae* strains identified 14 proteins whose electrophoretic mobility suggest they carry an additional charge, presumably due to acetylation (Takakura et al., 1992). Two other studies used two-dimensional difference gel electrophoresis (2D-DIGE) combined with mass spectrometric analysis to identify acetylated ribosomal proteins. In these studies, 30 and 19 of the 68 and 60 identified ribosomal proteins were N^α^-terminally acetylated, and 24 and 17 of these were NatA substrates, only 4 and 2 were NatB substrates and two were unusual NatD substrates, respectively (Arnold et al., 1999; Kamita et al., 2011). This indicates that a fair amount of ribosomal proteins are NTA; however, the functional consequences are not well understood.

A recent study suggests that NatA-dependent acetylation is necessary for ribosome biogenesis in conjunction with Ebp2. The nuclear protein Ebp2 is required for ribosome synthesis, especially for the maturation of 25S rRNA and assembly of the 60S subunit. Mutation of nuclear protein Ebp2 (*ebp2-14* mutant) exhibited synergistic growth defects and defects in the biogenesis of the 60S ribosomal subunit with *NAA10*Δ or *NAA15*Δ mutants, which could be repressed by overexpression of ribosomal protein eL36A (Wan et al., 2013). Indeed, eL36 was found to be N-terminally acetylated by NatA in one of the previous screens (Arnold et al., 1999). However, mutation of the second amino acid to proline in eL36A or eL36B did not exhibit the above mentioned effects, whereas mutation of Brx1 (*brx1-S2P*), a functional partner of Ebp2, led to synthetic lethality with *ebp2-14,* suggesting that Brx1 and not eL36A acetylation by NatA is important for ribosome biogenesis (Wan et al., 2013). According to their sequence, all analyzed proteins in this study are putative NatA substrates. However, further studies are necessary to identify the exact acetylation targets that impact ribosome biogenesis to elucidate the mechanism by which NTA affects this process.

To analyze the impact of N^α^-terminal acetylation on translational activity, Kamita and colleagues performed a polyU-dependent poly-(Phe) synthesis assay with purified ribosomes from wild type and *NAA15*Δ strains (Kamita et al., 2011). The authors detected a decrease of the ribosomal protein synthesis activity by 27% in the mutant compared to wild type ribosomes (Kamita et al., 2011). In dilution spot assays, increased sensitivity of the *NAA15*Δ mutant cells towards the antibiotic paromomycin and hygromycin was observed and since both antibiotics induce errors during translation, the authors speculate that NTA of ribosomal proteins by NatA may be required to maintain proper translational fidelity (Kamita et al., 2011). Indeed, elevated stop codon readthrough was observed in the *NAA15*Δ mutant strain in a translational fidelity assay using a bicistronic reporter gene, and this effect was strongly influenced by the addition of paromomycin, indicating that N^α^-acetylation by NatA is required for optimal translational termination (Kamita et al., 2011). The authors claim to not find increased sensitivity of the *NAA15*Δ cells towards anisomycin and cycloheximide, which are known to affect peptidyl transferase activity and the translocation step in protein synthesis, respectively, and therefore speculate that NTA does not influence transferase activity or translational elongation (Kamita et al., 2011). However, the corresponding figure seems to show an increased sensitivity of the mutant cells towards both antibiotics. Similar experiments with *NAA10*Δ/*NAA15*Δ cells also showed an increased sensitivity of the double knockout towards cycloheximide (Van Damme et al., 2014), indicating that NTA by Naa10 might affect ribosomal functions on different levels. Since the above mentioned affects could also be indirect, possibly through acetylation of regulatory factors, further studies are necessary to analyze the exact functional and structural changes induced by NTA of ribosomal subunits.

In addition to the above, NatA-dependent acetylation could influence ribosome function through the action of ribosome-associated protein biogenesis factors (RPBs), which act on newly synthesized proteins as soon as they emerge from the ribosome exit tunnel. These RPBs regulate enzymatic processing, chaperone-assisted folding, and protein sorting. RPBs include the NATs, methionine aminopeptidases, signal recognition particle (SRP), as well as chaperones like the Ssb/Ssz/Zuotin triad and nascent polypeptide-associated complex (NAC). For a review, see (Kramer et al., 2009). Recent findings suggest that NatA regulates protein import into the endoplasmic reticulum (ER). In eukaryotes, this translocation process is mediated by the heterotrimeric Sec61 complex that consists of the yeast proteins Sec61, Sbh1 and Sss1. During co-translational translocation, the nascent polypeptide chain is recognized by the signal recognition particle SRP that, through its interaction with the SRP receptor, targets the ribosome nascent chain complex (RNC) to the membrane of the ER. The elongating polypeptide chain then is transferred to Sec61 and inserted into the channel. In contrast to this, posttranslational translocation requires, besides the Sec61 complex, also the Sec63 complex (Sec63, Sec62, Sec71 and Sec72) and the luminal chaperone BiP. Additionally, in yeast, another complex, called Ssh1, can be formed by Ssh1, Sss1 and Sbh2 that is exclusively responsible for co-translational import into the ER. For recent reviews, the reader is referred to (Robson and Collinson, 2006; Park and Rapoport, 2012; Akopian et al., 2013).

Some data indicates that NatA preferentially associates with translating ribosomes but might not bind to at least some ribosomes translating certain SRP substrates. The sequential steps of this process are not really known. Particularly, Naa15 was enriched in the polysomal fractions from extracts rich in randomly translating ribosomes as compared to a ribosomal fraction from extracts of non-translating ribosomes (generated by glucose deprivation) as shown by quantitative immunoblotting (Raue et al., 2007). To assess the binding properties of ribosome associated factors to ribosome nascent chain complexes (RNCs) carrying specific nascent polypeptides, the authors fused the nascent polypeptides of Pgk1 (monomeric soluble protein localized to the cytosol), ppα-factor (precursor that matures into the secreted pheromone α-factor) and Dap2 (type II membrane protein that is finally localized to the vacuole) to an N-terminal FLAG and isolated the specific RNCs from *in vitro* translation reactions (see also Table 2). In these experiments, Naa15 and Ssb1 could be cross-linked to purified RNCs loaded with Pgk1 and ppα-factor, whereas SRP was exclusively found to be cross-linked to RNC-Dap2 (Raue et al., 2007). In line with this, systematic bioinformatic studies of predicted N-terminal processing found that cytoplasmic proteins are typically biased in favor of acetylation, whereas secretory proteins are not (Forte et al., 2011). A differential N-terminal combined fractional diagonal chromatography (COFRADIC) study on cytosolic and organellar fractions of A-431 cells revealed that the degree of NTA is lower for integral membrane proteins compared to their soluble counterparts, supporting this idea (Aksnes et al., 2015c). Furthermore, mutation of the penultimate amino acid of the post-transcriptionally targeted proteins CPY (carboxypeptidase Y), Pdi1 and ppα-factor that allowed acetylation by NatA inhibited their translocation, whereas this effect was restored in *NAA10*Δ cells (Forte et al., 2011). Thus, the authors conclude that N^α^-acetylation is a general preclusion for efficient ER-targeting. However, a trafficking defect observed on artificially acetylated proteins does not provide sufficient proof of a general inhibitory role for NTA in ER-targeting, since these proteins are not NTA *in vivo*. The artificial SRP-substrates OPY and D_HC_-α translocated normally and were found to be unprocessed, even after mutation of the N-terminus (Forte et al., 2011). In this context it is noteworthy that both SRP and NatA bind to the ribosome via uL23/uL29 at the tunnel exit (Pool et al., 2002; Halic et al., 2004; Polevoda et al., 2008). Therefore, the authors argue that SRP could compete with NatA for binding to the ribosome thereby preventing NTA of the nascent polypeptide chain (Forte et al., 2011). In contrast to this, Soromani and colleagues found that two Sec proteins, Sec61 and Sec62, which are themselves targeted to the ER membrane in an SRP-dependent fashion, are acetylated by NatA (Soromani et al., 2012), indicating that NatA-dependent acetylation is still possible even if SRP is binding to specific RNCs. Similarly, the above mentioned proteome-wide study showed that NTA is more frequent than anticipated, occurring on 82% of all transmembrane proteins identified, although to a lesser extent, when compared to cytosolic proteins (Aksnes et al., 2015c). One explanation for the non-exclusive action of SRP and NatA might be related to the identification of additional SRP-binding sites on the ribosome. When SRP contacts the signal peptide and its receptor on the ER membrane, SRP is repositioned to the side opposite to uL23/uL29, freeing up uL23, allowing binding to the Sec61 translocon (Pool et al., 2002).

Taken together, this could indicate that NTA might interfere with post-transcriptional, SRP-independent protein translocation only but does not regulate cotranslational, SRP-dependent ER-targeting. However, different models suggest that SRP function is influenced by NAC and that this dynamic interplay is important for protein targeting to the ER (Zhang et al., 2012). Also, NAC has been shown to bind uL23 (Wegrzyn et al., 2006), although other studies identified eL31as the major ribosomal binding site of NAC (Pech et al., 2010; Zhang et al., 2012). Therefore, it would be interesting to see if and how NatA may be involved in this process and what the consequences are for the acetylation status of proteins. *NAA10*Δ cells showed no gross ER translocation defect for the ppα-factor, Sec62, or Pdi1 (Forte et al., 2011; Soromani et al., 2012).

As a side note, another Sec protein, Sbh1, is N-terminally acetylated by NatA, and because this protein is inserted by the guided-entry of tail-anchored protein insertion (GET) pathway, the authors conclude that this translocation pathway is also not affected by NTA (Soromani et al., 2012) (Table 2).

**Table 2:**
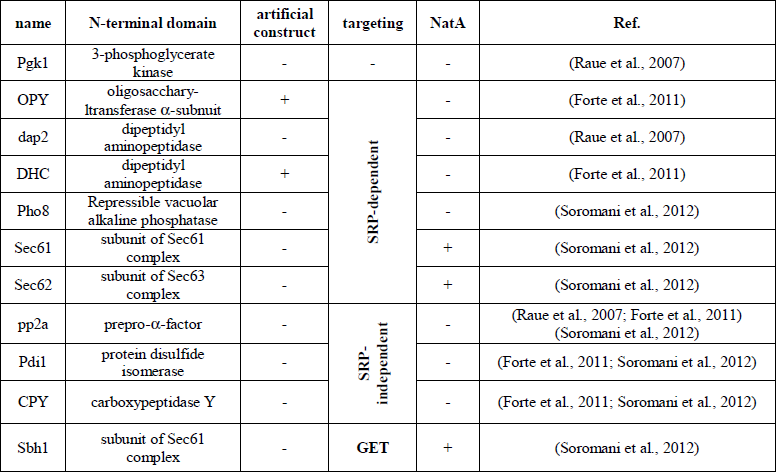
Summary of the proteins/artificial constructs used in the mentioned studies to analyze protein targeting. Pgk1 is a cytosolic protein and not targeted to the ribosome. Sbh1 is targeted by guided-entry of tail-anchored protein insertion (GET) pathway. SRP-dependent = co-translational targeting to the ER. SRP-independent = post-translational targeting

| name | N-terminal domain | artificial construct | targeting | NatA | Ref. |
| --- | --- | --- | --- | --- | --- |
| Pgk1 | 3-phosphoglycerate kinase | - | - | - | (Raue et al., 2007) |
| OPY | oligosaccharyltransferase α-subunit | + | SRP-dependent | - | (Forte et al., 2011) |
| dap2 | dipeptidyl aminopeptidase | - |  | - | (Raue et al., 2007) |
| DHC | dipeptidyl aminopeptidase | + |  | - | (Forte et al., 2011) |
| Pho8 | Repressible vacuolar alkaline phosphatase | - |  | - | (Soromani et al., 2012) |
| Sec61 | subunit of Sec61 complex | - |  | + | (Soromani et al., 2012) |
| Sec62 | subunit of Sec63 complex | - |  | + | (Soromani et al., 2012) |
| pp2a | prepro-α-factor | - | SRP-independent | - | (Raue et al., 2007; Forte et al., 2011)<br>(Soromani et al., 2012) |
| Pdi1 | protein disulfide isomerase | - |  | - | (Forte et al., 2011; Soromani et al., 2012) |
| CPY | carboxypeptidase Y | - |  | - | (Forte et al., 2011; Soromani et al., 2012) |
| Sbh1 | subunit of Sec61 complex | - | GET | + | (Soromani et al., 2012) |

#### 3.2.3 NatA and proteasome function

The proteasome is an ATP-dependent protease consisting of a cylinder-shaped 20S core flanked by a 19S regulatory subcomplex. The regulatory particle is organized in a lid (regulatory particle non ATPase Rpn3, -5, -9, -11, -12, -15) and a base, comprised of the organizing subunits Rpn1 and Rpn2, the ubiquitin receptors Rpn12 and Rpn13, as well as 6 AAA-ATPases (regulatory particle triple A protein1-6, Rpt1-6), that unfolds the respective substrate, opens the gate and translocates the substrates into the 20S core. The 20S core consists of 4 stacked heptameric ring structures, two outer rings that are composed of 7 structural α subunits and 7 catalytic β subunits that exhibit protease activity in the center of the particle. For a recent review see (Bhattacharyya et al., 2014).

Some of the catalytic β-type subunits of the 20S core are synthesized as catalytically inactive forms with an N-terminal propeptide, which is then cleaved during particle assembly. Incomplete propeptide removal results in severe assembly defects of the proteasome (Groll et al., 1997). Mutants in β5, β2 and β1, which lack the N-terminal propeptide, are defective for specific peptidase activity, are more sensitive to environmental stresses, and have defects in proteasome assembly (Arendt and Hochstrasser, 1999). Deletion of *NAA10* or *NAA15* restores the phenotype, indicating that the propeptide protects the N-terminal threonine of the mature form against acetylation (Arendt and Hochstrasser, 1999). A similar study showed that the deletion of the N-terminal propeptides of the β1, β2 subunit precursors drastically inhibited the enzymatic activity of the proteasome (Jäger et al., 1999). Further chemical analyses revealed that acetylation of the N-terminal threonine in β1 is responsible for the measured activity loss, supporting the idea that the propeptide protects the mature subunit moiety from being acetylated prior to incorporation into the proteasome complex (Jäger et al., 1999). The authors also propose a similar mechanism for β2 and β5; however, the N-termini have not been analyzed for NTA (Jäger et al., 1999). Taken together, this data indicates that NTA of catalytic β-chains would interfere with their activity. The expression of these proteins together with a protective propeptide and subsequent cleavage during particle assembly could be one evolved mechanism to escape the potent acetylation machinery.

Furthermore, acetylation of other components of the proteasome has been identified *in vivo*. Purified 20S proteasome subunits from wild type, *NAA10*Δ*, NAA20*Δ and *Naa30*Δ mutant cells indicated that the α1, α2, α3, α4, α7, and β3 subunits were acetylated by NatA, the β4 subunit was acetylated by NatB, and the α5 and α6 subunits were acetylated by NatC (Kimura et al., 2000). In a similar approach for the 26S regulatory subunit, the same group found that 8 subunits, Rpt4, Rpt5, Rpt6, Rpn2, Rpn3, Rpn5, Rpn6, Rpn8, Rpn13 and Rpn15 were NatA substrates and that 2 subunits, Rpt3 and Rpn11, were NatB substrates, whereas Rpt1 was unprocessed (Kimura et al., 2003; Kikuchi et al., 2010). Although these data indicates that a multitude of proteasomal factors are acetylated *in vivo*, little is known about the functional consequences of this modification and the so far analyzed effects seem to be relatively small. Analysis of the 20S proteasomes derived from *NAA15*Δ mutant cells revealed a very similar *in vitro* activity compared to wild type-derived proteasomes with respect to chymotrypsin-like, trypsin-like, and peptidylglutamyl peptide hydrolyzing activities (Kimura et al., 2000). In the absence of the chaotropic agent SDS, which is used in the *in vitro* assay to induce activity of the 20S proteasome, the *NAA15*Δ proteasomes displayed a twofold-higher chymotrypsin-like peptidase activity (Kimura et al., 2000). However, crude proteasome extracts from the wild type strain and the *NAA15*Δ deletion mutant exposed a similar accumulation level of the 26S proteasome *in vitro* and exhibited normal chymotrypsin-like activity in the presence of ATP (Kimura et al., 2003). On the other hand, the accumulation level of 20S proteasome and the chymotrypsin-like activity were appreciably higher in the extracts from the *NAA15*Δ deletion mutant (Kimura et al., 2003), indicating that NTA might have a suppressing role in proteasome function.

In contrast to this, data generated with *H. volcanii* suggests that acetylation is required for the assembly of the 20S core particle. In this archaea, the structural proteins α1 and α2 are acetylated on their initiator methionine (Humbard et al., 2006). The acetylated form of α1 was almost exclusively found in purified S20 core particles, even after the cellular ratio of acetylated/non-acetylated α1 was changed by mutation of the penultimate amino acid, indicating that the acetylated protein is preferably inserted into the core particle (Humbard et al., 2009). However, the observed changes on methionine cleavage or NTA of α1 that were introduced by mutation of its penultimate residue did not follow the classical rules. Variants of α1 with Ser, Thr or Val and even Pro were found to be acetylated on the initiator methionine instead of the penultimate amino acid after methionine processing. The reason for this unexpected behavior is not clear.

Recently, the effect of NTA on nuclear-cytoplasmic shuttling of proteasomes during starvation was analyzed. In *S. cerevisiae*, proteasomes accumulate in the nucleus; however, during starvation, proteasomes are relocalized into the cytosol and stored into cytoplasmic proteasome storage granules (PSG). This shuttling efficiency ceases with age (impairment of proteasome relocalization and/or PSG formation correlates with replicative age). NatC auxiliary subunit Naa35 was identified in a yeast screen for mutants affecting nuclear-cytoplasmic shuttling during starvation (van Deventer et al., 2014). Additionally, loss of N-acetylation by deletion of the NatC subunits Naa30, Naa35 or Naa38 or a catalytically inactive NatC mutant (Naa30 N123A/Y130A) caused an increased nuclear enrichment/retention of the proteasomes under starvation without affecting PSG formation as visualized by tagging the catalytically active β1-subunit (Pre3) of the proteasome with GFP (van Deventer et al., 2014). Disruption of NatB (*NAA20*Δ and *NAA25*Δ) had similar effects but also quenched PSG formation under starvation, whereas disruption of NatA (*NAA10*Δ, *NAA15*Δ and *NAA50*Δ) had no effects (van Deventer et al., 2014). The associated NatB or NatC substrates have not been identified and mutation of the N-terminus of the putative candidates α5 (Pup2), α6 (Pre5), Rpn9, Fub1, Avo2, Hul5 or Nup100, that would prevent acetylation (MX- to MP-mutation), failed to phenotypically mimic cells lacking NatB or NatC.

Proteasomes from rat (Tokunaga et al., 1990) and humans have been found to be acetylated as well. Purification of human 26S proteasome from HEK293 cells and LC-MS/MS identified eight subunits, Rpt3, Rpt5, Rpn1, Rpn2, Gankyrin, α 5, α6, and β4, that are acetylated at the first methionine residue, whereas seven subunits, Rpt4, Rpt6, Rpn6, Rpn13/ADRM1, α4, α7, and β3 are acetylated at the second amino acid after the removal of the first methionine residue (Wang et al., 2007). It would be interesting to see whether NTA of the identified proteins has any influence on enzymatic activity or assembly of the proteasome in these organisms. Recently, experiments linked Naa10 function to the 28S proteasome in human cells. In the 28S proteasome, the 19S regulatory subcomplex is replaced by a 11S cap (proteasome activator 28; consisting of a heteroheptamer of PA28α and PA28β) that stimulates protein degradation in an ATP-independent manner without recognizing ubiquitin (Bhattacharyya et al., 2014). Knockdown or overexpression of Naa10 in RKO and H1299 cells increased or reduced chymotrypsin-like proteasome activity, respectively, without affecting the expression level of PA proteins, indicating that Naa10 is a negative regulator of 28S proteasome activity (Min et al., 2013). Experiments in a cell free system revealed that Naa10 reverts activating effects of PA28 on proteasome activity of 20S proteasome core but had no effect on chymotrypsin-like activity of the 20S core in the absence of PA28β or 26S proteasome (Min et al., 2013).

Taken together, from mass spectrometrical analyses it is known that NTA of proteasomal subunits by NatA, NatB and NatC is quite common in various organisms, however, the functional consequences of this modification are largely obscure. Recent efforts indicate that NTA could affect proteasomal assembly and might regulate catalytic activity, possibly dependent on stress situations such as cellular starvation. On the other hand, in case of the β-subunits, the N-terminal Thr of the mature form is indispensable for catalytic activity. Since acetylation would interfere with such an activity, these proteins are synthesized as inactive form containing a N-terminal propeptide which is cleaved during particle assembly.

#### 3.2.4 NatA in prion propagation

Several studies have linked NatA to the Sup35/[*PSI^+^*] prion in *S. cerevisiae*. Prions are protein-based units of inheritance that, by altering their conformational state and self-assembly into amyloid fibers, serve as novel, self-replicating agents to determine phenotypic traits (Sweeny and Shorter, 2008). The eukaryotic release factor 3, eRF3/Sup35 facilitates, in its soluble nonprion [*psi^-^*] state, translation termination. In the [*PSI^+^*] prion state, Sup35 adopts an alternative conformation that promotes self-aggregation, which subsequently leads to a termination defect (stop codon read-through) (Liebman and Chernoff, 2012). The N-terminal PrD (prion-determining domain) of Sup35 is required for formation and propagation of the prion state.

Disruption of *NAA10* or *NAA15* in [*PSI^+^*] yeast strains reversed the [*PSI+*] phenotype as measured by stop-codon readthrough assays (Pezza et al., 2009). Furthermore, Sup35 aggregates from NatAΔ [*PSI+*] strains exhibit a reduced thermodynamic stability and are composed of smaller core polymers, compared to wild type [*PSI+*], but the number of Sup35 aggregates or the solubility of Sup35 seemed to be unaltered (Pezza et al., 2009). However, later studies by the same group showed that the decrease in Sup35 aggregate size observed in a [*PSI+*]^strong^ NatAΔ strain leads to increased ribosome binding of the aggregates and a decreased stop-codon readthrough, indicating an increase in aggregate-associated termination activity (Pezza et al., 2014). Taken together, these data suggest that NatA function is required for [*PSI+*] strains to display the prion phenotype. Sup35 was identified as a substrate of NatA in a global qualitative and quantitative analysis of protein N-acetylation (Arnesen et al., 2009b) and mutation of the penultimate amino acid to proline abrogates NTA of Sup35. S2P mutation in [*PSI+*] induced a reduced stability and core size of Sup35-S2P aggregates; however, the effects were weaker compared to deletion of NatA, and the Sup35-S2P did not revert stop-codon readthrough, indicating that Sup35 NTA alone is not sufficient and additional factors contribute to the phenotype conversion of the NatAΔ [*PSI^+^*] strain (Holmes et al., 2014).

The propagation of the [*PSI*^+^] state involves the action of many chaperones and recent evidence indicates that acetylation of these proteins contributes to the prion phenotype. The Hsp104 chaperone and the Hsp70 chaperone Ssa1/2 (constitutively expressed) and Ssa3/4 (stress inducible), promote fragmentation of prion fibers into smaller seeds, initiating new rounds of prion propagation (Liebman and Chernoff, 2012). The Hsp40 protein Sis1 is an important co-chaperone of HSP70 in prion propagation (Higurashi et al., 2008; Tipton et al., 2008). In contrast to this, the Hsp70 chaperone Ssb1/2 seems to destabilize the prion state [*PSI^+^*] (Liebman and Chernoff, 2012). In an early study on acetylated proteins in yeast, Ssa1/2 and Ssb1/2 proteins but not Hsp104 have been shown to undergo NTA by NatA (Polevoda et al., 1999). Notably, Ssb was also found to be acetylated by ssNaa10 in *S. solfataricus*, despite the fact that the N-terminal sequence (MEEK…) of Ssb points towards NatB as the corresponding NAT (Mackay et al., 2007; Polevoda et al., 2009). This again indicates that the sequence specificity of the NAT has co-evolved with its substrates sequence. However, the sequence of Ssa3/4 (starting MS…) also indicates that these proteins are targets of NatA in yeast, which has been confirmed biochemically (Van Damme et al., 2014). Loss of NatA strongly increased heat-shock response (HSR) as measured by Hsf1 reporter gene expression, increased Hsp104, Ssa1/2, Ssb1/2 and Sis1 protein levels and induced a change in the recruitment of the chaperones Ssb1/2 Ssa1/2 to Sup35 as seen in immunoprecipitation experiments in [*PSI+*] cells (Holmes et al., 2014), but did not affect Hsp104 or Ssa1/2 expression levels in [*psi^-^*] cells (Pezza et al., 2009). Additionally, elevation of HSR activity by using a constitutively active Hsf1 mutant (Hsf1ON), reduces Sup35 aggregate size in [*PSI+*] cells (mimics the effects of NatA disruption), whereas downregulation of HSR by expressing a dominant-negative mutant of Hsf1 (Hsf1DN) resulted in the loss of the smallest Sup35 core polymers (Holmes et al., 2014). Because disruption of NatA in [*PSI+*] strain led to the accumulation of protein aggregates, as assayed by a centrifugation assay, sensitivity to translational inhibitors and Hsf1-mediated gene expression, the authors conclude that loss of NTA promotes general protein misfolding, a redeployment of chaperones to these substrates, and a corresponding stress response that subsequently reduces the size of prion aggregates and reverses their phenotypic consequences (Holmes et al., 2014). However, single mutation of Ssa1, Ssa2, Ssb1, Ssb2 or Sup35 only partially mimicked the effects of NatAΔ (Pezza et al., 2009; Holmes et al., 2014), whereas mutation of all of these proteins [*ssa1(S2P)*Δ*ssa2, ssb1(S2P)*Δ*ssb2, sup35(S2P)*] did mimic the effects of disruption of NatA in [*PSI+*] strain, including induction of smaller sized Sup35 aggregates and reversion of stop-codon readthrough in colony growth *ade1-14* (UGA) assays (Holmes et al., 2014). This indicates that increased chaperone protein levels and loss of Sup35 NTA combinatorially contribute to the effect of NatA disruption on prion propagation (Holmes et al., 2014).

### 3.3 NATs in prokaryotes (archaeabacteria, eubacteria)

In contrast to eukaryotes, in *Escherichia coli* and *Salmonella typhimurium,* only a few proteins seem to be N^α^-terminally acetylated (Bernal-Perez et al., 2012; Ansong et al., 2013) and ectopically expressed human proteins in *E. coli* usually lack N^α^-acetylation as shown for human ββ alcohol dehydrogenase (Höög et al., 1987), rat chaperonin 10 (Ball et al., 1994), Annexin II tetramer (Kang et al., 1997) or Spartin and Tropomyosin (Johnson et al., 2010). However, some proteins are found to be fully or partially acetylated when expressed in *E. coli*, including stathmin-like subdomains (Charbaut et al., 2002), prothymosin α (Wu et al., 2006), interferon-γ (Honda et al., 1989), interferon-α (Takao et al., 1987) and the serine proteinase inhibitor eglin c (Grütter et al., 1985). Also, a recent analysis of 112 million mass spectrometric datapoints from 57 bacterial species revealed that NTA might be markedly more common than previously known, as about 16% of bacterial N-termini were found to be N-acetylated in some species (Bonissone et al., 2013). Structural homology searches revealed that the structure of NATs is conserved from ancestral to prokaryotic and eukaryotic cells with homologous proteins in *S. pombe*, *H. sapiens*, *S. typhimurium*, *A. tumefaciens*, *S. mutans*, and *T. acidophilum* (Chang and Hsu, 2015).

To date only three N^α^-acetyltransferases have been identified in prokaryotes which specifically modify only one protein each, as opposed to eukaryotes, in which these enzymes are much less specific (Nesterchuk et al., 2011). These enzymes, RimI, RimJ and RimL (ribosomal modification) specifically acetylate the ribosomal proteins bS18 (Isono and Isono, 1980; Yoshikawa et al., 1987), uS5 (Cumberlidge and Isono, 1979; Yoshikawa et al., 1987) and bL12 (Isono and Isono, 1981; Tanaka et al., 1989), respectively. According to their low sequence identity with the NATs it is most likely that the RIM proteins do not have a common ancestor and evolved independently (Polevoda and Sherman, 2003a; Vetting et al., 2008).

Mutation of each of these Rims leads to temperature sensitivity in the *E. coli* strain K12 (Isono and Isono, 1978) and there is evidence that the RimI and RimJ-mediated acetylation occurs post translationally and is required for the assembly of the ribosomal subunits or translation initiation (or more generally for ribosome function) (Poot et al., 1997; Recht and Williamson, 2001; Roy-Chaudhuri et al., 2008).

RimJ has evolved dual functionality; it functions in ribosomal protein acetylation and as a ribosome assembly factor in *E. coli* (Roy-Chaudhuri et al., 2008). Furthermore it has been shown to regulate pyelonephritis-associated pilus (Pap) transcription in response to multiple environmental cues (White-Ziegler et al., 2002).

RimL seems to have a regulatory function during the growth cycle. A shift in acetylation occurs during the growth cycle: L7 is the acetylated form of L12 (Terhorst et al., 1973). L12/L7 is not acetylated during early log phase but becomes acetylated at the stationary phase (Ramagopal and Subramanian, 1974). Later mass spectrometric data from intact ribosomes showed that NTA of L12 during the stationary phase correlates with an increased ribosomal stability which mediates the adaption of the cells to nutrient deprivation (Gordiyenko et al., 2008).

Taken together, it seems like the main function of NTA in *E. coli* is regulation/initiation of translation/transcription. Whether acetylation can also act as a signal for protein degradation, as recently proposed for NTA in Eukarya (Hwang et al., 2010), has to be shown. However, this is questioned as the enzyme that recognizes acetylated N-termini in eukaryotes and tags them for destruction (Doa10 ubiquitin ligase) does not have orthologues in prokaryotes such as *E. coli* (Jones and O’Connor, 2011). Another puzzling fact is the recent finding that even in bacteria NTA is more common in some species than others (Bonissone et al., 2013). This would either indicate that the RIM proteins are either not as specific as previously anticipated or that another, yet unidentified, acetyltransferase is responsible for this modification.

#### 3.3.1 In Archaea

In contrast to bacteria, NTA is quite common in archaea. Large-scale proteomics studies in *Halobacterium salinarum*, *Natronomonas pharaonis* (Falb et al., 2006) and *Haloferax volcanii* (Kirkland et al., 2008) showed that up to 14%, 19% and 29% of the identified proteins are acetylated, respectively, although many of the encountered proteins were only acetylated partially. In the former study, acetylation was limited to cleaved N-termini starting with serine or alanine residues, suggesting NatA activity (Falb et al., 2006; Aivaliotis et al., 2007), whereas the latter study also found NatB like substrates (Kirkland et al., 2008).

So far, only one NAT in *Sulfolobus solfataricus* has been studied in more detail. However, homologous proteins have also been identified in the archaea *Pyrobaculum aerophilum*, *Natronomonas pharaonis* and *Halobacterium salinarium* (Mackay et al., 2007), *Aeropyrum pernix* (Polevoda and Sherman, 2003b) as well as *Thermoplasma acidophilum* (Han et al., 2006) and *Thermoplasma volcanium* (Ma et al., 2014). The *S. solfataricus* ssNaa10 represents an ancestral variant of the eukaryotic NATs and it adopts a more relaxed substrate specificity as it acetylates NatA, B, C and E substrates (Mackay et al., 2007). No homologs of Naa15 have been identified in *S. solfataricus*, suggesting that ssNaa10 is solely responsible for NTA in this species. In line with this, *in vitro* acetylation assays showed that, although ssNaa10 has higher sequence similarity to hNaa10 over hNaa50 (34% vs. 24% sequence identity, respectively) and therefore has a preference for Ser- over Met-amino-terminal substrates, it is still able to accommodate NatA and NatE substrates (Liszczak and Marmorstein, 2013). Furthermore, studies on the X-ray crystal structure of ssNaa10 combined with mutagenesis and kinetic analyses showed that the active site of ssNaa10 represents a hybrid of the NatA and NatE active sites (Liszczak and Marmorstein, 2013), explaining the above findings. A later study confirmed these findings and showed that Glu35 has a critical role for the unique substrate specificity of ssNaa10 (Chang and Hsu, 2015). The X-ray crystal structure of archaeal *T. volcanium* Naa10 has also been reported, revealing multiple distinct modes of acetyl-Co binding involving the loops between β4 and α3 including the P-loop (Ma et al., 2014). The acetylation activity of ssNaa10 has been confirmed *in vitro* for a variety of substrates including the *S. solfataricus* proteins Holliday junction resolving enzymes (Hjc and Hje), single-stranded DNA-binding protein (SSB), as well as Alba1 (Mackay et al., 2007; Ma et al., 2014).

Until now, no auxiliary subunit of any of the NATs has been described in archaea, and the function of NTA in archaea is unknown.

### 3.4 NATs in plants

In 2003, Polevoda and Sherman identified the first candidates for NATs (Naa10, Naa20 and Naa30) in *A. thaliana* by homology search with the amino acid sequences of yeast NATs (Polevoda and Sherman, 2003b). More recent studies identified orthologs and their gene duplicates for 6 catalytic subunits and 5 auxiliary subunits corresponding to NatA-F from humans in *Populus nigra* (Liu et al., 2013) and *Populus trichocarpa*, *Chlamydomonas reinhardtii*, *Medicago truncatula*, and *Vitis vinifera* (Bienvenut et al., 2012), implying that all known NATs are conserved in woody plants. Additionally, six distinct methionine aminopeptidases have been identified and characterized in *A. thaliana* (Giglione et al., 2000).

Accordingly, up to 1054 proteins in *A. thaliana* (Baerenfaller et al., 2008; Bienvenut et al., 2012) and 58 proteins in *Populus nigra* (Liu et al., 2013) have been found to be N-terminally acetylated to date. A large-scale N-terminomic study in the model diatom *Thalassiosira pseudonana* (phytoplankton) revealed that about 70% of cytosolic proteins were completely or partially acetylated (Huesgen et al., 2013). Furthermore, similar acetylation patterns between animal and plant kingdoms have been observed, suggesting a strong convergence of the characterized modification (Liu et al., 2013). Also, the observed patterns for methionine removal follow the same rules as those of other eukaryotes (Huesgen et al., 2013).

In addition to cytosolic proteins, posttranslational acetylation of proteins imported into organelles has been observed. A large scale analysis of chloroplast preparations from *Arabidopsis thaliana* identified 47 N-terminal acetylated nuclear encoded proteins (Zybailov et al., 2008) and proteomic studies detected NTA on 30% of nuclear encoded stromal chloroplast proteins in the green algae *Chlamydomonas reinhardti* (Bienvenut et al., 2011) and on 50% of nuclear encoded plastid proteins in *Thalassiosira pseudonana* (Huesgen et al., 2013). Similarly, a comparative large scale characterization of plant and mammalian proteomes found that 25 % of the identified acetylated *Arabidopsis thaliana* proteins become N^α^-acetylated at a position downstream of the annotated position 1 or 2 (compared to 8 % in humans) and the majority of these proteins (>80 %) reside in the chloroplasts and become acetylated at the new N-terminus after removal of the chloroplast transit peptide (Bienvenut et al., 2012). Furthermore, the authors indicate that the sequence of the acetylated chloroplast appears to match to NatA substrates, suggesting a dedicated NatA might exist in this organelle (Bienvenut et al., 2012). In *Thalassiosira pseudonana,* a nuclear encoded plastid-targeted putative N-acetyltransferase was identified that could be responsible for the observed NTA of imported and plastid-encoded proteins (Huesgen et al., 2013).

All these data indicate that NTA is very common in plants. Also, it appears that most NAT components have been identified in the analyzed species so far, and the fact that the same rules for N-terminal processing apply indicates that N-terminal processing is very well conserved in plants. Plants also possess chloroplasts and many studies support post-translational NTA in these plastids. It would be very interesting to identify and characterize the dedicated acetyltransferase in these plastids to understand the evolution of the NATs.

#### 3.4.1 Function of NTA in plants

Compared to mammals or yeast, the function of NTA in plant is even less studied and understood. However, some data points out that NTA in plants may have similar effects on protein function and stability as in eukaryotes. The ε-subunit of chloroplast ATP synthase has been found to be partially acetylated in *Citrullus lanatus* (wild watermelon), and during drought, the non-acetylated form gets degraded by metalloaminopeptidases (Hoshiyasu et al., 2013), indicating that the effect of NTA on stability might be more complex and might even depend on the respective substrate. However, in contrast to the proposed destabilizing Ac/N-end rule pathway in the mammalian cytosol (Varshavsky, 2011), NTA may have a stabilizing role in the plastids of plants. For example, analysis of the acetylation status of stromal proteins in combination with pulse-chase experiments revealed that among all identified N-terminally acetylated stromal proteins with a short half-life, NTA was underrepresented (Bienvenut et al., 2011). The authors speculate that an α-acetylated N-terminus could protect stromal proteins from rapid degradation. They base this speculation on a) the majority of the plastid proteins have an unusual N-terminus due to the cleavage of their transit peptides and b) the chloroplast does not have a proteasome which is indispensable for the ^Ac^N-degron degradation process, but contains a number of bacterial-type chambered proteases, including ClpP (which appears to display specificity for such unusual N-termini), (Bienvenut et al., 2011). However, further studies have to analyze the effect of NTA on the stability of plastid and cytosolic plant proteins.

It should be mentioned that NTA might be generally important for photosynthesis. An *Arabidopsis thaliana* Naa30 (*At*Mak3) mutant exhibits a decreased effective quantum yield of photosystem II and a growth defect (Pesaresi et al., 2003). However, in the latter case, it is not clear how a cytoplasmic N-acetylation defect would influence chloroplast function. The authors speculate that NTA may be relevant for “stability and/or import of organellar precursor proteins and thus, indirectly, for organellar function” (Pesaresi et al., 2003). This could indicate that, as discussed for yeast, targeting of proteins is regulated by NTA. In this context it is interesting to mention that components of the photosystem II also have been found to be acetylated in *Spinacia oleracea* (spinach) (Michel et al., 1988). Thus, direct acetylation of photosystem compounds by an organelle-localized NAT could also influence photosystem function.

As for mammalian NATs, NTA has also been implicated in plant development. Systematic investigation of publicly available microarray data showed that the expression levels of NatA-F are relatively low during development; these NATs are expressed at low levels but share distinct tissue-specific expression patterns (Zhu et al., 2014). Furthermore, a loss of function mutation of Naa20 (NatB catalytic subunit, nbc-1) or Naa25 (NatB auxiliary subunit, transcurvata2, tcu2) in *Arabidopsis thaliana* caused a strong phenotype that includes leaf reticulation, early flowering, unfertilized or aborted ovules in siliques, indicating that the NatB auxiliary subunit is important for vegetative and reproductive development (Ferrández-Ayela et al., 2013). Genetic interaction of *NAA25* and *AGO10* (Argonaute10) suggest a link between NatB-mediated N^α^-terminal acetylation and the microRNA pathway (Ferrández-Ayela et al., 2013). The mechanism for this genetic interaction is not clear and future studies have to explore whether other NATs are also involved in plant development.

## 4 Open Questions

NTA is one of the most abundant protein modifications known, and the NAT machinery is very well conserved throughout all Eukarya. Over the past 50 years, the function of NTA has begun to be slowly elucidated, and this includes the modulation of protein-protein interaction, protein-stability, protein function and protein targeting to specific cellular compartments. Many of these functions have been studied in the context of Naa10/NatA; however, we are only starting to really understand the full complexity of this picture. Roughly, about 40 % of all human proteins are substrates of Naa10 and the impact of this modification has only been studied for a few of them. However, recent publications have linked mutations in Naa10 to various diseases, emphasizing the importance of Naa10 research in humans. Besides acting as a NAT in the NatA complex, recently other functions have been linked to Naa10, including post-translational NTA, lysine acetylation and NAT/KAT-independent functions.

### Co-translational NTA and protein quality control

The highly complicated process of protein translation requires the interplay of multiple factors, including non-ribosomal proteins that permanently or transiently associate with the ribosome and welcome the emerging protein as soon as a nascent polypeptide reaches the exit from the ribosomal tunnel. This “welcoming committee” consists of ribosome-associated protein biogenesis factors (RPBs) that co-translationally regulate various processes, including protein folding, targeting protein sorting, protein quality control and protein modifications (Giglione et al., 2014). RPBs include the NATs, MetAPs (methionine aminopeptidases), SRP (signal recognition particle) and chaperones like Hsp70/Hsp40 and NAC (nascent polypeptide-associated complex) (Kramer et al., 2009). The region around the exit tunnel of the proteasome, which is comprised of the ribosomal proteins uL22, uL23, uL24, uL29 as well as eL19, eL31 and eL39 in archaea and eukaryotes, constitutes a general docking platform for RPBs, and recent research indicates that uL23 might play an eminent role as it interacts with almost all RPBs analyzed (Kramer et al., 2009).

It is widely accepted that NATs and NatA, in particular, are linked to the ribosome and co-translationally acetylate nascent chain as they emerge from the ribosomal exit tunnel (Strous et al., 1973; Filner and Marcus, 1974; Strous et al., 1974; Driessen et al., 1985; Yamada and Bradshaw, 1991; Gautschi et al., 2003; Arnesen et al., 2005a; Raue et al., 2007; Polevoda et al., 2008; Arnesen et al., 2009a; Arnesen et al., 2010). Recent findings suggest that NatA negatively regulates protein sorting/targeting to the ER through SRP-dependent and/or SRP-independent mechanisms, and the bias observed in secretory proteins towards not being NTA supports this idea, although some ER-targeted proteins, including Sec61 and Sec62, have been shown to be NTA by NatA (Raue et al., 2007; Forte et al., 2011; Soromani et al., 2012). Both, SRP and NatA bind to the ribosome via uL23/uL29 at the tunnel exit; however, SRP seems to reposition to the opposite side of the tunnel during signal peptide recognition to allow binding of the Sec61 translocon to uL23 (Pool et al., 2002; Halic et al., 2004; Polevoda et al., 2008). In addition to SRP and NatA, NAC has been shown to interact with uL23; however, eL31 seems to be the major ribosomal binding site of NAC (Wegrzyn et al., 2006; Pech et al., 2010; Zhang et al., 2012). This data on the one hand indicates that RPBs compete for nascent chains at the ribosome as suggested for NATs and N-myristoyltransferases (Utsumi et al., 2001) and on the other hand suggests that NATs might functionally regulate adjacent RPBs. We are only beginning to understand the complicated interplay of RPBs on the ribosome, but recent studies suggest that NAC might modulate the function of SRP possibly through the Hsp70/40 chaperone system in an isoform-specific fashion, and this dynamic interplay is important for proper protein targeting and folding (Koplin et al., 2010; del Alamo et al., 2011; Sedwick, 2011; Zhang et al., 2012; Pechmann et al., 2013; Holmes et al., 2014).

The role of Naa10/NatA in this context is obscure but the data summarized here strongly implicates a connection of NATs to the protein sorting and protein folding machinery, which could explain, at least in part, the connection of Naa10 and other NATs to neurodegenerative diseases such as Alzheimer’s or Parkinson’s disease, where misfolding of certain protein species leads to the accumulation of toxic amyloid aggregates (Asaumi et al., 2005; Zabrocki et al., 2008; Pezza et al., 2009; Kang et al., 2012; Kang et al., 2013; Holmes et al., 2014; Pezza et al., 2014). However, the ribosome itself is susceptible to acetylation by NatA and other NATs (Takakura et al., 1992; Arnold et al., 1999; Kamita et al., 2011), and NTA of ribosomal proteins seem to affect various processes, including ribosome biogenesis, ribosomal fidelity/translational activity and translational termination (Kamita et al., 2011; Wan et al., 2013).

The challenge of future studies would be to analyze the dynamic interplay of NatA and other NATs with the ribosome, the nascent polypeptide chain and its associated factors such as SRP, NAC and other chaperones. Structural studies of larger ribosomal complexes could help gain insight into the molecular events occurring directly during the birth of new proteins. Recent advances in the Cryo-EM field have already led to high-resolution structures of the ribosome, nascent polypeptide chain, and the Sec61 complex (Voorhees et al., 2014) or nascent chain-containing 60S-Listerin-NEMF complex (Shao et al., 2015). Similarly, new techniques such as proximity-specific ribosome profiling (Jan et al., 2014) could be used to monitor the location of the NAT machinery on translating ribosomes, which would enhance the knowledge of co-translational NTA in general and elucidate the mechanistic interplay between NATs and other ribosome-associated protein biogenesis factors.

Also, most of what we know about the co-translational NTA process is derived from data generated in *S. cerevisiae*. It would be interesting to see whether these concepts are conserved in higher eukaryotes.

### Post-translational acetylation by Naa10

Besides acting co-translationally, accumulating reports have reported post-translational NTA by Naa10. Indeed, despite being reported to be mainly associated with ribosomes in yeast, a major pool of hNaa10 exists in a ribosome-free context, and a fraction of cytosolic hNaa10 even exists independent of the NatA complex (Arnesen et al., 2005a; Van Damme et al., 2011b). Similarly, NatB has also been shown to not always co-express and co-localize with its catalytic counterpart Naa20 in differentiated neurons in mouse, implying that the auxiliary subunit may function either with an unidentified NAT protein partner(s) or possibly in a NAT-independent manner (Ohyama et al., 2012). However, large-scale N-terminomic analyses are shedding light on post-translational NTA by identifying N-terminally acetylated proteins starting beyond position 1 or 2, such as mitochondrial or chloroplast proteins in plants, that become post-translationally NTA after cleavage of their transit peptide (Zybailov et al., 2008; Helbig et al., 2010; Bienvenut et al., 2011; Helsens et al., 2011; Van Damme et al., 2011b; Bienvenut et al., 2012; Huesgen et al., 2013; Van Damme et al., 2014). With respect to acetylation, at least for lysine, in mitochondria, it is hypothesized to occur chemically without any enzyme due to the high Ac-CoA concentration in this organelle. In any case, the prerequisite for amino acetylation is the deprotonation of the amine that allows subsequent attack on the carbonyl carbon of Ac-CoA. Since an unperturbed ε-amino group (pK_a_ = 10.4) is almost exclusively in the protonated state at physiological pH, non-catalytic acetylation is only possible when the local environment depresses the pK_a_ values into near neutral range (e.g. pK_a_ = 8.4) by virtue of nearby positive charges or by desolvation (Ghanta et al., 2013). The pK_a_ of a N^α^-terminal amino-group is even lower (pK_a_ =8) and therefore non-enzymatic NTA could be possible at high Ac-CoA concentrations; however, to our knowledge no study has addressed chemical N^α^-terminal acetylation in mitochondria.

Until now, it is not clear if a dedicated NAT enzyme resides in organelles or if Naa10 is imported into the respective organelle (Bienvenut et al., 2012; Van Damme et al., 2014). In the latter case, the occurrence of classical NatA-substrates in organelles would suggest that somehow the complete NatA complex is present in organelles, since monomeric Naa10 seems to exhibit altered substrate specificity, preferentially acetylating acidic N-termini. The reason for this change are structural rearrangements of the substrate binding pocket in Naa10 induced by Naa15 binding (Liszczak et al., 2013). In general, acidic N-termini are not very common since methionine aminiopeptidases do not cleave the iMet if the penultimate amino acid is Asp, Glu, Asn or Gln (Xiao et al., 2010). Acidic N-termini can either be directly generated by proteolytic cleavage or, by enzymatic deamidation of exposed Asn or Gln through the N-terminal asparagine amidohydrolase (NTAN1) and N-terminal glutamine amidohydrolase (NTAQ1), respectively (Sriram et al., 2011). In the case of γ- and β-actin, the only known non-canonical Naa10 substrates, the iMet is first acetylated by NatB, which is then believed to be cleaved by N^α^-acetylaminopeptidase to expose the acidic N-terminus (Sheff and Rubenstein, 1992; Polevoda and Sherman, 2003b; Van Damme et al., 2011b; Van Damme et al., 2014). According to the N-end rule, acidic N-termini are further modified by arginyl-transferases and targeted for degradation (Varshavsky, 2011). One example in this case is IAP (inhibitor of apoptosis). Caspase-dependent cleavage of IAP generates a Asn-bearing N-degron and degradation by the Arg/N-end rule pathway is indispensable for regulating apoptosis (Ditzel et al., 2003). Since both monomeric Naa10 and arginyl-transferases act on acidic N-termini, a crosstalk of NTA and the Arg/N-End rule is possible; however, that remains to be proven.

In conclusion, despite the profound accomplishments in the field regarding co-translational NTA by Naa10, the significance of post-translational NTA is still very inconclusive. The main proof for post-translational NTA to date is the detection of acetylated internal peptides, although this finding does not exclude co-translational effects. However, since the nuclear localization of Naa10 is widely accepted, a post-translational acetylation of nuclear substrates could also be conceivable, but a detailed study of NTA proteins in the nucleus is still missing. *In vitro* acetylation studies in combination with structural investigations suggest a substrate switching of uncomplexed Naa10. The biological consequences of this have not yet been studied in detail, but could lead to several possibilities, including: 1) there are more Naa10 substrates than anticipated; 2) these non-canonical substrates are mainly products of proteolytic cleavage; and 3) these substrates are also targeted by arginyl-transferases, suggesting a competition with the Arg/N-End rule pathway. Similarly, despite the fact that ε-acetylation of Naa10-substrates has been shown by many biochemical studies, KAT-activity is still very controversial in the field, since structural studies argue against the possibility of lysine chains being inserted into the catalytic center.

### NAT-independent functions of Naa10 and transcriptional regulation

Many studies have addressed the intercellular localization of Naa10 in a variety of cell systems, showing that Naa10 mainly localized to the cytoplasm and to a lesser extent in the nucleus, but also isoform specific localization patterns of Naa10 have been suggested (Fluge et al., 2002; Sugiura et al., 2003; Bilton et al., 2005; Arnesen et al., 2006a; Arnesen et al., 2006b; Chun et al., 2007; Xu et al., 2012; Park et al., 2014; Zeng et al., 2014). In agreement with this, a functional nuclear localization signal (KRSHRR) has been identified and characterized (Arnesen et al., 2005a; Park et al., 2012; Park et al., 2014). Naa15 also harbors a putative NLS; however, analysis of the nuclear localization of Naa15 revealed discrepant results (Willis et al., 2002; Arnesen et al., 2005a).

The function for a possible nuclear redistribution of Naa10 is not yet clear, but accumulating studies showed that Naa10 directly or indirectly regulates transcriptional activity in different pathways, including Wnt/β-catenin, MAPK, JAK/STAT5, AP-1, mTOR, NF-kB and BMP signaling (Lim et al., 2006; Kaidi et al., 2007; Lim et al., 2008; Kuo et al., 2009; Kuo et al., 2010; Seo et al., 2010; Park et al., 2012; Xu et al., 2012; Yoon et al., 2014; Zeng et al., 2014). The exact mechanism for this is not known and, despite speculations that Naa10 might directly act as a transcription factor, to our knowledge, a direct interaction of Naa10 and promoter regions has only been described for the E-cadherin promoter (Lee et al., 2010a) and the c-Myc promoter (Lim et al., 2008).

Despite the fact that the NAT-dependent function of Naa10 has been addressed in numerous studies, a possible NAT-independent function of Naa10 is widely understudied and the mechanism of such action is still not clear. The use of an enzymatic-dead hNaa10 mutant helped to differentiate between NAT-dependent and -independent functions. This R82A mutation is located in a R^82^-x-x-G^85^-x-A^87^ consensus sequence, critical for the binding of acetyl-CoA, and has been shown to exhibit low acetyltransferase activity and inhibited autoacetylation of Naa10 (Asaumi et al., 2005; Seo et al., 2010; Seo et al., 2014). According to the widely accepted idea that the acetyltransferase activity of Naa10 is crucial for its function, disruption of the catalytic activity abrogated Naa10 function to regulate APP/Aβ40 endocytosis and secretion, mTOR signaling, NF-κB activation, cyclin D1 expression and HIF-1α stability (Asaumi et al., 2005; Kuo et al., 2010; Seo et al., 2010; Park et al., 2012; Seo et al., 2014). However, in some cases, eliminating NAT activity did not affect Naa10 function. For example, reintroduction of Naa10 R82A restored colony forming in Naa10-depleted cells, although not to the same degree as wild type hNaa10, and overexpression of the mutant increased DNMT activity similarly to wild type Naa10 (Lee et al., 2010a). The authors conclude that Naa10 facilitates DNMT1-mediated gene silencing in an acetyltransferase (AT)-independent manner by recruiting DNMT1 to the E-cadherin promoter, where DNMT1 can silence the transcription of the tumor suppressor E-cadherin through methylation (Lee et al., 2010a). Similarly, Naa10 has been shown to competitively bind to PIX proteins in an AT-independent manner, thereby disrupting the GIT-PIX-Paxillin complex, resulting in reduced activation of the cell migration machinery (Hua et al., 2011). However, the authors did not completely rule out acetylation-dependent mechanisms in regulating cell mobility, as the Naa10-R82A mutant may still have some remaining function to acetylate some other specific proteins that participate in cell mobility control (Hua et al., 2011). A different study showed that Naa10 regulates Janus kinase 2-STAT5a transcriptional activity independent of its acetyltransferase activity (Zeng et al., 2014). Particularly, overexpression of either wild type Naa10 or Naa10-R82A decreased STAT5a binding to the ID1 promoter, inhibited ID1 protein levels and reduced migration to a similar level in MCF-7 and MDA-MB-231 breast cancer cells (Zeng et al., 2014). Furthermore, because STAT5a was not found to be acetylated at lysine residue(s) and its protein expression was not affected by the exogenous wild-type or mutant Naa10, the authors speculate that the acetyltransferase activity of Naa10 could be dispensable for inhibiting STAT5a-dependent *ID1* expression and suppressing invasiveness of breast cancer cells (Zeng et al., 2014). Although the sequence of STAT5a (starting: MAG…) matches the substrate specificity of Naa10, the authors did not report the NTA status of STAT5a. In another study, AT-independent activity Naa10 has been implicated in the activation of the 28S proteasome; however, the mechanism for this action is not clear (Min et al., 2013). The authors speculate that Naa10, through binding to components of the 28S activation complex PA28, could sterically hinder peptide entrance into the proteasome, although this would need further investigation (Min et al., 2013).

The goal of future studies will be to analyze whether Naa10 indeed acts as a transcription factor and/or modulates transcription indirectly through acetylation of substrates or via other NAT-independent functions. The recent design and synthesis of the first 3 bisubstrate inhibitors that potently and selectively inhibit the NatA/Naa10 complex, monomeric Naa10, and hNaa50 (Foyn et al., 2013a), further increases the toolset to discriminate between NAT-dependent and independent function. Additionally, it would be interesting to analyze the mechanism and the signal pathways involved in the nuclear import or a potential nucleo-cytoplasmic shuttling of Naa10.

## Acknowledgements

This review and the corresponding Gene Wiki article are written as part of the Gene Wiki Review series – a series resulting from a collaboration between the journal GENE and the Gene Wiki Initiative. The Gene Wiki Initiative is supported by National Institutes of Health (GM083924). Additional support for Gene Wiki Reviews is provided by Elsevier, the publisher of GENE. The laboratory of G.J.L. is supported by funds from the Stanley Institute for Cognitive Genomics at Cold Spring Harbor Laboratory (CSHL). The authors thank Jason O’Rawe for critical reading of the manuscript.

The corresponding Gene Wiki entries for this review can be found here: http://en.wikipedia.org/wiki/N-alpha-acetyltransferase_10 and here http://en.wikipedia.org/wiki/NAA15

## Competing Interests

G.J.L serves on the medical advisory board of GenePeeks, Inc. and the scientific advisory board of Omicia, Inc. The study did not involve these companies and did not use products from these companies.

